# Activation of PP2A-B56α leads to aberrant EGFR signaling and proliferative phenotypes in PDAC

**DOI:** 10.1101/2025.07.18.665598

**Authors:** Claire M. Pfeffer, Sydney J. Clifford, Saadia A. Karim, Ella Rose D. Chianis, Brittany N. Heil, Lauren E. Gartenhaus, Emma F. Kay, Garima Baral, Elizabeth G. Hoffman, Sagar M. Utturkar, Harish Kothandaraman, Whitney Smith-Kinnaman, Kasi Hansen, Emily G. Smith, Elisabeth Porter, Fuping Zhang, Nadia Atallah Lanman, Jukka Westermarck, Emma H. Doud, Amber L. Mosley, Jennifer P. Morton, Brittany L. Allen-Petersen

**Author notes:** Corresponding Author Brittany L. Allen-Petersen, Purdue University. 201 S. University St, West Lafayette, IN 47907.

## Abstract

Pancreatic ductal adenocarcinoma (PDAC) stands to become the second most deadly cancer by 2030. The small GTPase, KRAS, is mutated in over 90% of PDAC patients and considered the primary driver mutation. Despite being an almost ubiquitous event, KRAS mutations have been difficult to target therapeutically, particularly KRAS^G12D^, the most common mutation in PDAC. In addition to these pharmacological challenges, KRAS mutations have been shown to drive signaling plasticity and therapeutic resistance through phosphorylation cascades in most cancers. Protein phosphatases are master regulators of kinase signaling, however the contribution of phosphatase deregulation to mutant KRAS cancer phenotypes is poorly understood. Protein phosphatase 2A (PP2A) inhibits effectors downstream of KRAS, placing this family of enzymes as key regulators of PDAC oncogenic signaling. However, our previous studies utilizing small molecule activating compounds of PP2A show a heterogeneous response in PDAC, with some cell lines displaying increased oncogenic signaling despite induction of phosphatase activity. Similarly, specific PP2A subunits exhibit both tumor suppressive and oncogenic functions depending on the cellular context. Therefore, understanding the role of PP2A in regulating cancer phenotypes is critical for the future development of therapeutic strategies that leverage this phosphatase. Here, we determined the impact of the specific PP2A subunit, B56α, on PDAC phenotypes using both genetic and pharmacological activation strategies in human PDAC cell lines and genetic mouse models. We demonstrate that while PP2A-B56α suppresses specific oncogenic pathways, B56α activation exacerbates PDAC proliferative phenotypes and decreases overall survival *in vivo*, potentially through increased epidermal growth factor receptor (EGFR) signaling. EGFR is a critical signaling node in PDAC as inhibition or loss of EGFR prevents KRAS-driven tumorigenesis and increased EGFR activity is associated with poor patient outcome. The activation of EGFR by PP2A-B56α is in part mediated through increased expression and processing of EGFR ligands, specifically amphiregulin, heparin-binding EGF-like growth factor (HB-EGF), and epiregulin. Furthermore, pharmacological PP2A activation in combination with EGFR inhibitors mitigates this signaling and increases cell death. Together, these studies implicate a previously undescribed non-canonical role for PP2A-B56α in EGFR signaling that contributes to PDAC progression.

## INTRODUCTION

Pancreatic cancer is the 3^rd^ leading cause of cancer-related deaths in the United States and has a five-year survival rate of just 13%^1^. Over 90% of pancreatic ductal adenocarcinoma (PDAC) patients have driver mutations in the small GTPase, KRAS^2^. The aberrant activation of KRAS signaling has been shown to alter a wide range of signaling cascades leading to highly aggressive and therapeutically refractory disease^2–4^. As there are limited therapeutic options for PDAC patients, the identification of signaling mechanisms that contribute to or regulate KRAS-driven PDAC is of the utmost importance.

Protein phosphatases are master regulators of cellular signaling cascades but their role in cancer remains poorly understood. Protein phosphatase 2A (PP2A) is a large family of Ser/Thr phosphatases that negatively regulate many oncogenic pathways including KRAS^5–7^. PP2A is a heterotrimeric holoenzyme which is composed of an A (scaffolding) subunit, B (regulatory) subunit, and C (catalytic) subunit. Therefore, the function and targets of PP2A vary depending on which specific B subunit is incorporated into the complex. There are four distinct PP2A B subunit families (B/B55, B’/B56, B’’/PR72, B’’’/STRN) with multiple genes and splice variants in each family. While there is some functional redundancy amongst the B subunits, the impact of individual B subunits is context specific. The B56α subunit has been implicated to have tumor suppressor function with loss of B56α driving tumorigenic phenotypes^8,9^. *In vivo*, we have shown that the loss of B56α accelerates the formation of KRAS^G12D^-driven PDAC precursor lesions suggesting that B56α plays a critical role in suppressing PDAC initiation^10^. Based on these findings, we hypothesize that activation of PP2A-B56α may negatively regulate PDAC-associated oncogenic pathways and attenuate PDAC tumor progression. In support of this hypothesis, pharmacological activation of PP2A decreases tumor burden in multiple preclinical models, including mutant KRAS lung and pancreatic cancer^11–17^. The small molecule activator of PP2A (SMAP), DT-061, globally activates PP2A but displays a preferential stabilization of B56α-containing heterotrimeric PP2A complexes^18^.

Here, we sought to understand the role of PP2A-B561? in regulating late-stage PDAC progression. In contrast to the tumor suppressive roles of B561?, we have identified a novel feedback mechanism in a subset of PDAC cell lines in which PP2A-B56α activation drives Epidermal Growth Factor Receptor (EGFR) signaling through upregulation of EGFR ligands. This activation exacerbates proliferation in PDAC cell lines which can be attenuated with loss of EGFR. Similarly, the conditional loss of the PP2A B56 inhibitor, Cip2a, in an aggressive PDAC mouse model significantly decreases overall survival and drives increased EGFR phosphorylation. Finally, the combination of PP2A activators and EGFR inhibition is synergistic *in vitro* and *in vivo*. Together these findings support a potentially adverse feedback loop through EGFR in response to PP2A-B56α activation in PDAC.□□□

## RESULTS

### Loss of B56α reduces proliferation in PDAC cell lines

As B56α has been implicated as a B subunit contributing to PP2A’s tumor suppressive functions, we generated B56α knockout PDAC cell lines (AsPc1, MiaPaCa2, and Panc1) and assessed the impact on cell proliferation. Cells were first transduced and selected for stable expression of Cas9. Single cell Cas9 clones were then transduced with two independent CRISPR guide RNAs (sgB56α1 and sgB56α2) or a non-targeting (sgNT) guide. Single cell clones from selected cells were then screened for B56α loss by western blot (Figure 1A) and proliferation and clonogenic colony formation were then evaluated. Unexpectedly, loss of B56α led to a significant reduction in proliferation over 96 hours compared to sgNT control (Figure 1A). Similarly, we found that B56α loss resulted in a reduced ability to grow in a clonogenic colony formation assay relative to sgNT control (Figure 1B). Together, these results suggest that the loss of B56α attenuates the proliferative capacity of PDAC cell lines. Given B56α’s known tumor suppressive roles, these findings highlight a potentially contradictory role for B56α in PDAC.

**Figure 1.**
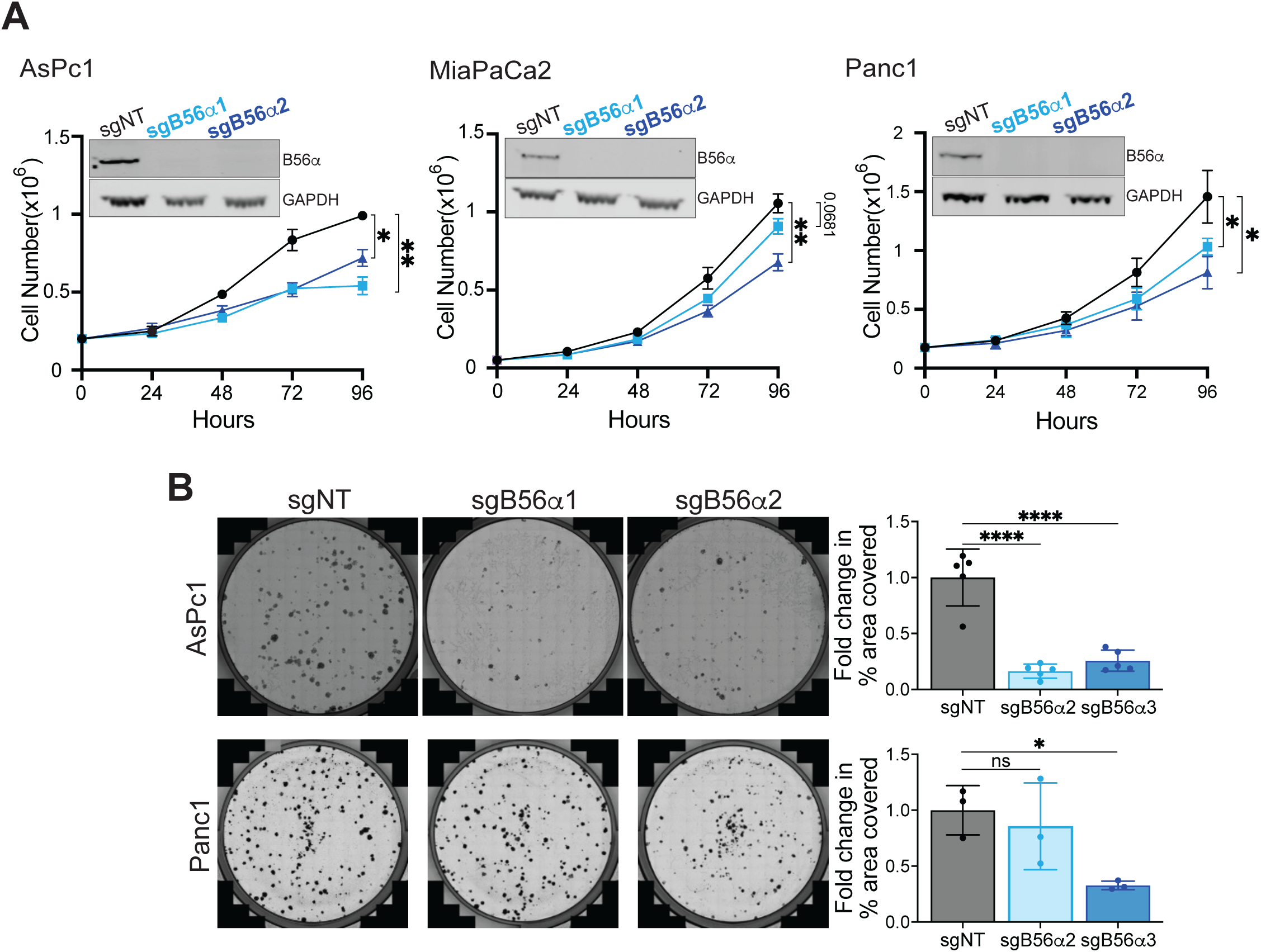
PDAC cell lines with loss of B56α have reduced proliferation. (A) Inset depicts representative western blot from at least 3 biological replicates from AsPc1, MiaPaca2, and Panc1 cells showing loss of B56α protein in B56α knockout cell lines (sgB56α1 and sgB56α2) compared to control non-targeting guides (sgNT). GAPDH was used as a loading control. Graphs depict proliferation (cell number) across 96 hours in the above cell lines comparing B56α knockout lines (sgB56α1/2) to non-targeting control cells (sgNT). Area under the curve was calculated from 3 biological replicates and a one-way ANOVA was run on the area under the curve to determine significance. (B) Representative images of clonogenic colony growth and quantification of percent area of the well covered in in AsPc1 and Panc1 cells comparing loss of B56α (sgB56α1/2) to non-targeting control cells (sgNT). Fold change was calculated from at least 3 biological replicates relative to sgNT and dots represent individual values for each biological replicate. Significance was determined using a one-way ANOVA. Error bars represent standard deviation *p<0.05, **p<0.01, ****p<0.0001

### Overexpression of B56α increases tumorigenic phenotypes in PDAC cell lines

Given that loss of B56α reduced proliferation, we then wanted to determine if increased B56α expression would result in the converse phenotype. AsPc1, MiaPaCa2, and Panc1 cell lines with stable overexpression of B56α (B56αOE) or empty vector (EV) were generated and B56α expression was confirmed by RT-qPCR and western blot (Supplemental Figure 1A, B). In contrast to B56α knockout phenotypes, B56αOE PDAC cells displayed a significant increase in proliferation, clonogenic colony growth, and soft agar colony growth relative to the EV control (Figure 2A-C). Given that there is functional redundancy amongst the PP2A B subunits, we asked if overexpression of the other subunits would recapitulate B56α overexpression phenotypes. To this end, we generated stable overexpression cell lines in AsPc1 with three B56 family subunits (B56γ, B56δ, and B56ε) as well as B55α (Supplemental Figure 1C, D). In contrast to B56α, the overexpression of B55α, B56γ, B56δ, or B56ε did not significantly alter proliferation or clonogenic colony growth compared to EV controls, suggesting that these phenotypes are specific to B56α in PDAC (Figure 2D, E). Finally, to determine if the increased tumorigenic phenotypes with B56α overexpression leads to similar tumor phenotypes in cancers driven by similar mutations, we created stable B56α overexpression lines in KRAS-mutant non-small cell lung cancer (NSCLC) cell lines (Supplemental Figure 1E). In the three NSCLC lines assayed (A549, H23, Calu6), overexpression of B56α failed to increase proliferation relative to EV control (Supplemental Figure 1F), suggesting that cancer cells do not respond uniformly to B56α expression. Together, these results contrast the known tumor suppressive roles of PP2A-B56α, and potentially points to a unique role for B56α in late-stage PDAC.

**Figure 2.**
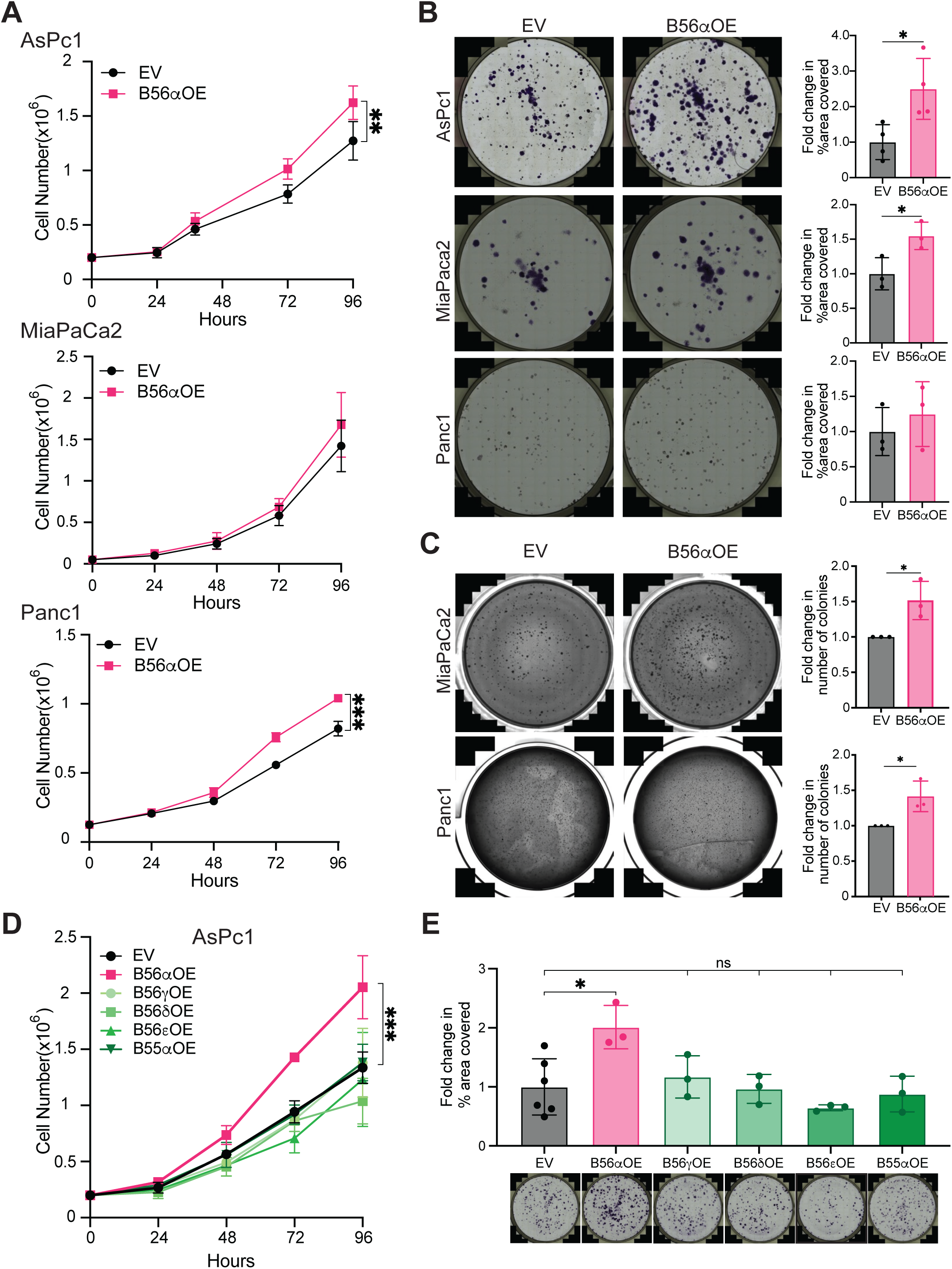
Overexpression of B56α increases proliferative phenotypes. (A) Proliferation across 96 hours in AsPc1, MiaPaca2, and Panc1 with overexpression of B561Z (B561ZOE) compared to empty vector cells (EV). Area under the curve was calculated from 3 biological replicates and two-tailed unpaired student’s t test was run on the area under the curve to determine significance. (B) Representative images of clonogenic colony growth in AsPc1, Panc1, and MiaPaca2 with B56αOE after 14 days. Fold change was calculated from at least 3 biological replicates (dots on graph) relative to EV. Significance was determined using a two-tailed unpaired student’s t test. (C) Representative images of growth in soft agar after 14 days in Panc1, and MiaPaca2 with overexpression of B56α (B56αOE). Fold change in number of colonies was calculated across 3 biological replicates (dots on graph) relative to EV control and two-tailed unpaired student’s t test was used to determine significance. (D) Proliferation over 96 hours in AsPc1 cells with overexpression of B56γ (B56γOE), B56δ (B56δOE), B56ε (B56εOE), B55α (B55αOE), or B56α (B56αOE) compared to EV control. Average values across 3 biological replicates were used to calculate area under the curve and a one-way ANOVA on the area under the curve was used to determine significance. (E) Representative images and quantification of clonogenic colony growth from cells shown in panel D. The average fold change of percent of well covered was calculated relative to EV control across 3 biological replicates and one-way ANOVA was used to calculate significance. Error bars represent standard deviation. Significance: *p<0.05, **p<0.01, ***p<0.001

### PP2A-B56α activation enriches for cell cycle and EGFR signaling

To identify the impact of PP2A activation on transcriptional programs, we treated a panel of PDAC cell lines (AsPc1, Panc1, and HPAFII) with DT-061 for 24 hours and performed RNAseq (Supplemental Tables 1-3). PP2A-B56α is a direct negative regulator of the oncoprotein c-MYC (MYC), with PP2A-B56α-driven dephosphorylation of MYC at S62 resulting in MYC degradation^17,19,20^. Therefore, as a read-out of PP2A activity, we analyzed the expression of MYC target genes using gene set enrichment analysis (GSEA)^21–23^. Within the genes that become downregulated in response to DT-061, there was a significant enrichment of MYC target genes across all three cell lines, consistent with PP2A activation (Supplemental Figure 2A). In contrast to the downregulation of MYC target genes, there were several datasets that were significantly enriched within genes that become upregulated in response to DT-061 treatment (Supplemental Table 4). Of the top 20 significantly enriched gene sets, seven were common amongst the three cell lines, including genes that are upregulated in response to EGFR activation (EGFR_UP.V1_UP), suggesting that PP2A activation may exacerbate EGFR signaling (Figure 3A). EGFR expression is known to increase during PDAC initiation and progression, and is overexpressed in PDAC compared to normal healthy pancreas^24,25^. Although EGFR is upstream of constitutively active mutant KRAS, loss of EGFR in pancreas-specific mutant KRAS PDAC mouse models (KRAS^G12D^ and KRAS^G12V^) prevents pancreatic tumorigenesis^24,25^, implicating EGFR signaling as a critical signaling node in PDAC. Furthermore, EGFR overexpression occurs in 40-90% of patients^25–27^ and correlates with poor patient survival highlighting the importance of EGFR signaling in PDAC^28,29^. DT-061 preferentially stabilizes PP2A complexes that contain the specific B subunit, B56α^30^. Therefore, to confirm that the enrichment of EGFR transcriptional programs is due to B56α signaling, HPAFII cells were transfected with an siRNA pool to B56α (siB56α) or non-targeting siRNA (siNT) and subsequently treated with DT-061. Knockdown of B56α prevented the DT-061-driven increase in EGFR activation genes, suggesting that the induction of EGFR signaling predominately functions through PP2A-B56α (Figure 3B, Supplemental Table 5, 6).

**Figure 3.**
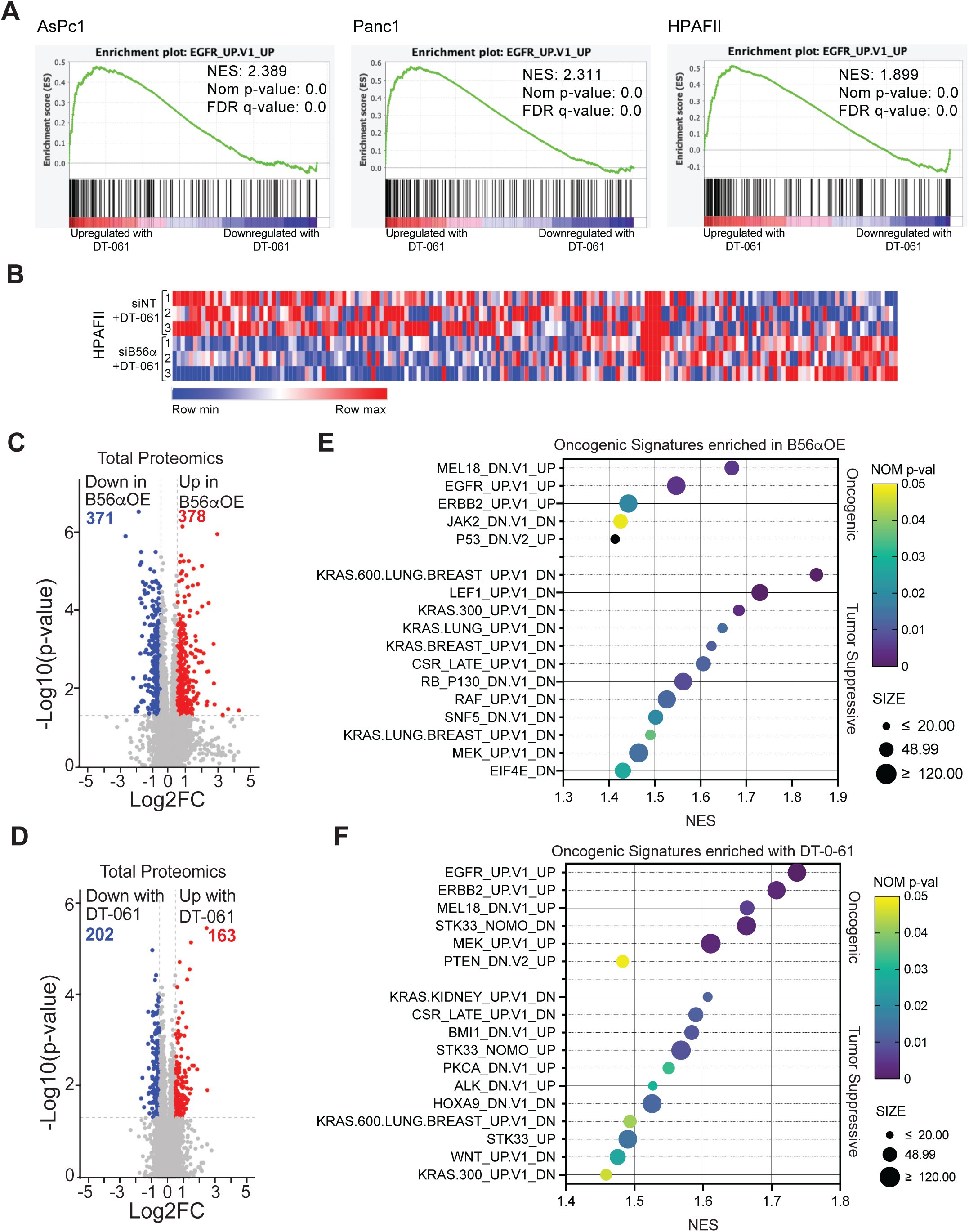
B56α overexpression or pharmacological activation of PP2A increases EGFR signaling. (A) GSEA analysis of RNAseq from AsPc1, Panc1, and HPAFII cells treated in technical triplicate with DT-061 showed a significant enrichment for genes associated with activated EGFR signaling compared to vehicle. (B) HPAFII cells with knockdown of B56α (siB56α) or non-targeting siRNA (siNT) were treated in technical triplicate with DT-061. Heat map of mRNA expression of genes within the EGFR gene set depicted in panel A. Volcano plot of differentially expressed total proteins from (C) AsPc1 cells with B56αOE compared to EV cells or (D) AsPc1 cells treated with DT-061 compared to vehicle. Cells were plated in technical triplicate. Significant proteins were defined as having a Log2 fold change (FC) above 0.5 and a -Log10 p-value of 1.3. Enriched oncogenic signature data sets from GSEA within the significant differential total proteins identified from (E) AsPc1 cells with B56αOE compared to EV cells or (F) AsPc1 cells treated with DT-061 compared to vehicle. Sets are separated by oncogenic function (top) and tumor suppressive function (bottom). Gene sets are arranged by normalized enrichment score (NES) and colored according to nominal p-value (NOM p-val). Size represents the number of proteins found in each gene set.

As pharmacological activation of PP2A altered EGFR transcriptional programs, we then performed proteomics and phospho-proteomics on AsPc1 EV and B56αOE cells, as well as AsPc1 cells treated with DT-061 or vehicle. In accordance with the role of PP2A as a phosphatase, DT-061 treatment predominately decreased phosphorylation with 306 phospho-sites significantly downregulated compared to 38 upregulated phospho-sites (Supplemental Figure 2B, Supplemental Table 7). Similar to previous reports, proteins with decreased phosphorylation in response to DT-061 were enriched for MYC targets, mitotic spindle, and mRNA processing^11,31,32^ (Supplemental Figure 2C). Similarly, overexpression of B56α resulted in the downregulation of 280 phospho-sites with similar enrichment (Supplemental Figure 2D, E). In contrast, 141 phospho-sites were upregulated in B56αOE cells. Consistent with the increased proliferation observed in B56αOE cells, proteins with increased phosphorylation were enriched for cell cycle programs and targets of cyclin dependent kinases (CDKs)(Supplemental Figure 2F, G).

Analysis of total proteomic changes identified that overexpression of B56α resulted in a significant downregulation of 371 proteins and upregulation of 378 proteins, while DT-061 treatment resulted in a significant downregulation of 202 proteins and upregulation of 163 proteins (Figure 3C, D, Supplemental Table 8). Similar to our RNAseq results, activation/overexpression of B56α resulted in a significant negative enrichment of MYC and E2F datasets (Supplemental Figure 2H,I). However, gene set enrichment analysis of oncogenic signature gene sets (C6) revealed a number of pathways upregulated in response to PP2A activation (Figure 3E,F). Within the top 17 significant datasets, five were found to be enriched in both B56αOE and DT-061 conditions, including EGFR and ERBB2 activation signatures. Together, these results suggest that while activation of PP2A-B56α is able to suppress known PP2A regulated pathways such as MYC, it also aberrantly increases EGFR signaling.

### PP2A-B56α increases EGFR ligand expression and availability to drive EGFR signaling

In PDAC, EGFR is rarely mutated and instead is commonly amplified or upregulated^29,33^. Therefore, EGFR ligands play a critical role in the activation of EGFR signaling during PDAC progression. Highlighting the necessity of ligand-driven EGFR signaling, loss of A Disintegrin and Metalloprotease 17 (ADAM17), which is required for processing five of the seven EGFR ligands, prevents mutant KRAS pancreatic tumorigenesis^25^. Furthermore, pancreata from KRAS mutant mice have increased mRNA expression of the EGFR ligands, transforming growth factor alpha (TGFα) and amphiregulin (AREG), relative to wildtype mice^25^. Given the critical roles of EGFR ligands in activating EGFR in PDAC, we asked if activation of PP2A-B56α increased EGFR ligand availability. AsPc1 cells were treated with DT-061 or vehicle for 6 hours, and the media was subsequently changed to remove the unbound drug and allowed to condition for 48 hours. Conditioned media was then filtered, added to AsPc1 parental cells for 24 hours, and then cell lysates were analyzed via western blot (Figure 4A). Cells treated with conditioned media from DT-061 treated cells had increased levels of phosphorylation at EGFR Tyr1068 (Y1068), a critical posttranslational modification that mediates downstream signaling, relative to vehicle conditioned media (Figure 4B, C). Similar results were obtained with conditioned media derived from AsPc1 B56αOE or EV cells added to AsPc1 EV cells for 1.5 hours (Figure 4D-F). Together, these findings indicate that both pharmacological and genetic activation of PP2A-B56α results in increased soluble factors capable of activating EGFR. To identify which EGFR ligands may contribute to PP2A activation, a panel of PDAC cell lines was treated with DT-061 for 12 hours and mRNA expression of EGFR ligands was measured. While DT-061 increased the expression of several EGFR ligands, AREG, epiregulin (EREG), and Heparin Binding EGF-like Growth Factor (HB-EGF) displayed the greatest fold increase relative to vehicle treated controls (Figure 4G).

**Figure 4.**
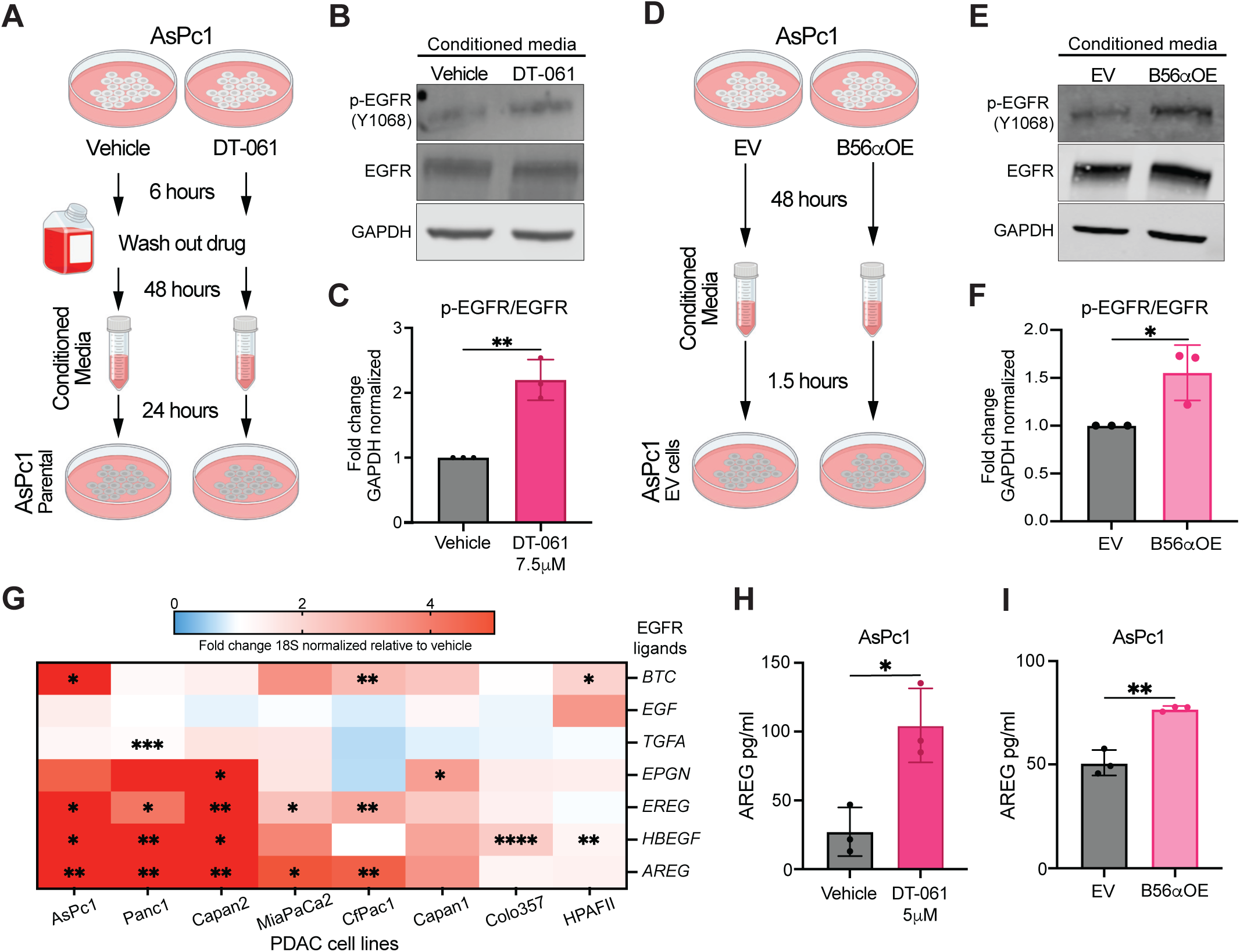
PP2A-B56α activation increases ligands which can activate EGFR. (A) Schematic for conditioned media (CM) experiment with DT-061 treatment. (B) Representative western blot of phosphorylated (Y1068) and total EGFR from AsPc1 cells treated with conditioned media (CM) from AsPc1 cells treated with 7.5μM DT-061 compared to vehicle CM. (C) Quantification of panel B. Fold change in phosphorylated EGFR relative to total EGFR normalized to GAPDH. Dots represent individual biological replicates. (D) Schematic for CM experiment with B561ZOE or EV cells. (E) Representative western blot of phosphorylated (Y1068) and total EGFR from AsPc1 cells treated with CM from AsPc1 cells with B56αOE or EV. (F) Quantification of panel E. Fold change in phosphorylated EGFR relative to total EGFR normalized to GAPDH. (G) mRNA expression of EGFR ligands from human PDAC cells lines plated in DMEM + 2% FBS and treated with 10μM DT-061 or vehicle (DMSO) for 12 hours. Average fold change was calculated relative to 18S expression and the vehicle condition for across biological replicates for each cell line. (H) ELISA quantification of processed AREG from the media derived from AsPc1 cells (H) treated with 5μM DT-061 or vehicle for 24 hours or (I) with B56αOE or EV. Dots represent individual biological replicates. Significance *p<0.05, **p<0.01, ***p<0.001, ****p<0.0001 by two-tail unpaired student’s t test.

Of all the EGFR ligands, AREG displayed the greatest increase in mRNA expression in response to PP2A activation. Increased expression of AREG has been shown to correlate with poor patient outcome, aggressive tumor phenotypes, and therapeutic resistance across several cancer types ^34–36^. In PDAC, AREG expression is increased in primary pancreatic tumors relative to healthy tissue, and in KRAS mutant PDAC mouse models compared to wild-type mice^25,37^. Additionally, AREG has been proposed as a diagnostic marker for distinguishing if pancreatic cysts are benign or malignant^38^. Furthermore, it has been recently identified that AREG signaling in PDAC cancer associated fibroblasts (CAFs) promotes a premetastatic environment^39^. Therefore, to identify if PP2A activation results in increased AREG shedding, AREG was quantified by ELISA from media derived from PDAC cells with either B56αOE or treated with DT-061. In AsPc1 and MiaPaCa2 cells, both genetic overexpression and pharmacological activation of PP2A resulted in a significant increase in AREG shedding (Figure 4H, I; Supplemental Figure 3A, B). In contrast, PP2A activation was unable to alter AREG shedding in either Capan2 or HPAFII cells (Supplemental Figure 3C-E), suggesting that there are subsets of PDAC cell lines that are more sensitive to alterations in PP2A signaling. Concurrently, EGF, HB-EGF, and EREG ELISAs were performed on the same conditioned media; however, all three ligands were below detectable values (<6.25pg/ml EREG, <3.9pg/ml EGF, and <16pg/ml HB-EGF). Together, these data suggest that PP2A-B56α activation induces AREG expression and processing.

### Increased proliferation in response to B56α overexpression is driven by AREG-EGFR signaling

Our data suggest that B56α overexpression in PDAC cells increases proliferation potentially through the activation of the AREG-EGFR signaling axis (Figure 1-4). To determine if EGFR is necessary for the B56α-dependent increase in PDAC proliferation, EGFR was knocked down using siRNA (Supplemental Figure 4A, B), and proliferation and migration were assessed. Over the course of 96 hours, knockdown of EGFR had no effect on the proliferation rates of EV cells (EV siEGFR vs EV)(Figure 5A). In contrast, EGFR inhibition was able to prevent the increased proliferation rates seen in B56αOE AsPc1 cells (B56αOE siEGFR vs B56αOE), with no significant difference observed between B56αOE siEGFR and EV cells (Figure 5A). Similar results were observed in a transwell migration assay, where B56αOE increased the migratory capacity of AsPc1 cells, which was significantly attenuated with knockdown of EGFR relative to siNT controls (Figure 5B, C). These data suggest that the B56α overexpression mediates tumorigenic phenotypes in part through EGFR signaling. Given that AREG shedding was increased with B56α overexpression (Figure 4), we then assessed the contribution of AREG to B56αOE proliferation rates. Similar to siEGFR, knockdown of AREG (siAREG) had no significant effect on AsPC1 EV cells. However, the suppression of AREG prevented the increased proliferation seen in B56αOE cells, returning proliferation rates back to baseline EV levels (Figure 5D, Supplemental Figure 4C). Knockout of B56α reduced PDAC proliferation (Figure 1). Therefore, we determined if the addition of exogenous AREG could restore the growth of the B56α knockout cells. Previous studies determined that 300pg/ml AREG potentially represents a diagnostic threshold for malignant PDAC cysts^38^. The addition of 300pg/ml AREG restored proliferation of AsPc1 B56α knockout cells back to levels seen in sgNT control conditions but had no effect on EV cells (Figure 5E). Together, these data suggest that the AREG-EGFR axis functions as a critical signaling pathway underlying the increased tumorigenic phenotypes that occur in response to B56α overexpression.

**Figure 5.**
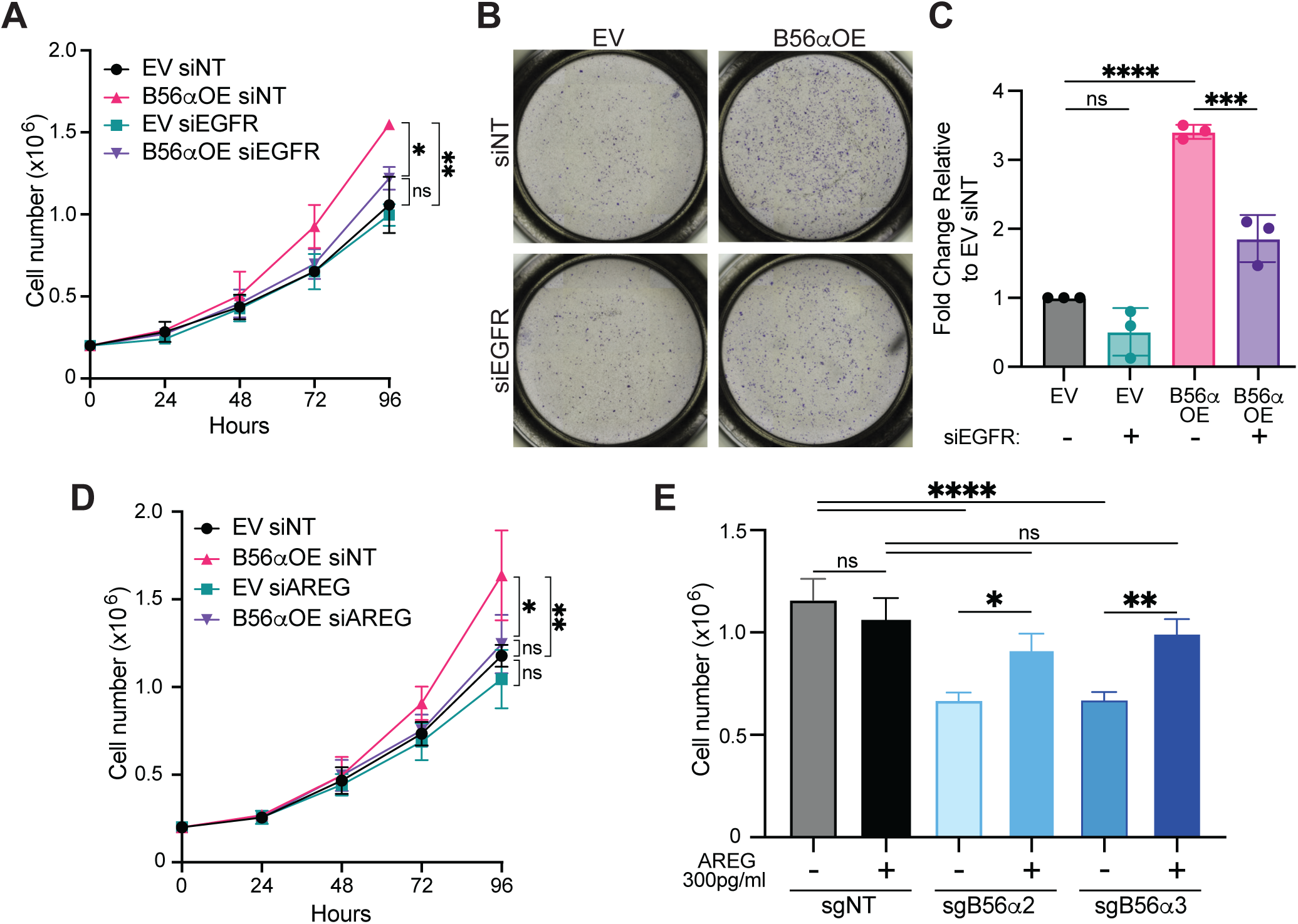
AREG and EGFR are necessary for increased proliferation with B56αOE. (A) Proliferation (cell number) across 96 hours in AsPc1 B56αOE compared to empty vector (EV) with either knockdown of EGFR (siEGFR) or non-targeting controls (siNT). Area under the curve was calculated from 3 individual biological replicates and one-way ANOVA was run on the area under the curve to determine significance. (B) Representative images of transwell migration of AsPc1 B56αOE cells compared to EV cells with knockdown of EGFR (siEGFR) or siNT. (C) Quantification of panel B. Average fold change in the number of migrated cells across biological replicates (dots) relative to EV siNT. (D) Proliferation across 96 hours in AsPc1 B56αOE compared to EV with knockdown of AREG (siAREG) or siNT. Area under the curve was calculated from 3 biological replicates and one-way ANOVA was run on the area under the curve to determine significance. (E) 300pg/ml AREG added to B56α knockout cells (shB56α1/2) rescues proliferation (cell number) at 96 hours post plating. Graph represents the average of 3 independent biological replicates. Significance *p<0.05, **p<0.01, ***p<0.001, ****p<0.0001 by one-way ANOVA.

### Genetic loss of the PP2A inhibitor, CIP2A, reduces overall survival in a model of PDAC *in vivo*

Previous studies have determined that knockdown of Cancerous inhibitor of PP2A (CIP2A), an endogenous cellular inhibitor of the PP2A B56 family^40^, decreases PDAC tumor growth and MYC signaling in Panc1 and Capan1 cells^41^. These results are consistent with our findings that PP2A activation suppresses MYC transcriptional programs (Figure 3). However, given that B56αOE cells also display an increase in EGFR signaling, we asked whether the chronic activation of PP2A-B56 in response to CIP2A loss would alter PDAC progression *in vivo*. For these studies, we generated a conditional *Cip2a* knockout mouse model of PDAC by crossing *Pdx1*-*Cre*, *Kras^LSL-G12D^*, *Tp53^LSL-R172H^* (KPC) mice with *Cip2a^fl/fl^* (Cip2a^fl/fl^) mice to generate KPCC^fl/fl^ (*Pdx1*-*Cre*, *Kras^LSL-G12D^*, *Tp53^LSL-R172H^*, *Cip2a^fl/fl^*) mice. KPC mice are considered the gold standard genetically engineered mouse model for PDAC, with KPC mice recapitulating both metastatic tumor progression and desmoplastic microenvironment found in PDAC patient tumors^42–44^. The genetic loss of this potent PP2A-B56 inhibitor accelerated PDAC tumorigenesis leading to a significant decrease in overall survival, with a median KPC survival of 148 days compared to 105 days in KPCC^fl/fl^ mice (Figure 6A). Furthermore, consistent with our *in vitro* findings, there was a significant increase in phosphorylated Y1068 EGFR in KPCC^fl/fl^ pancreatic tissues compared to KPC (Figure 6B, C). These results suggest that the chronic activation of PP2A exacerbates PDAC progression potentially through elevated EGFR signaling.

**Figure 6.**
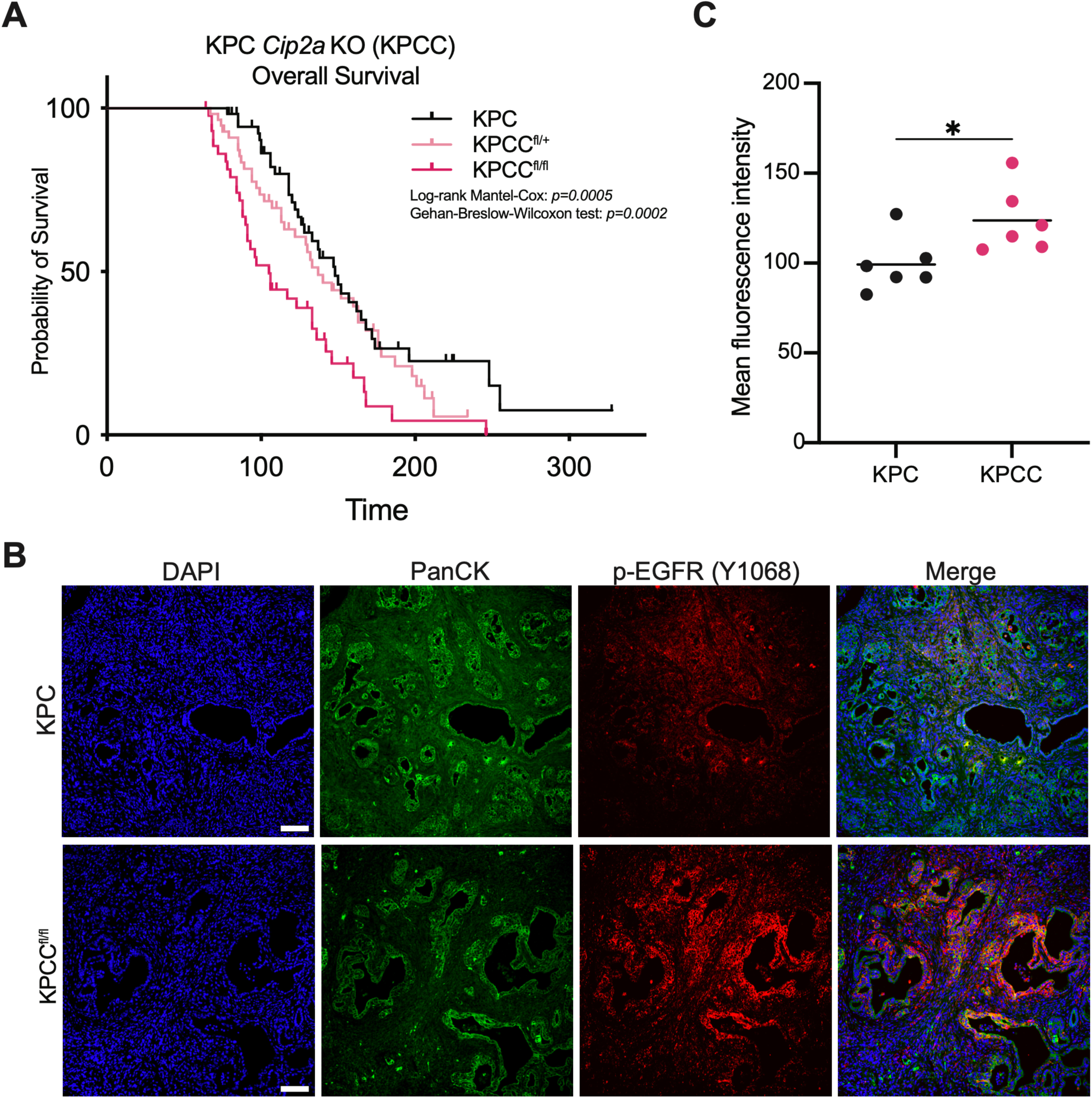
Loss of *Cip2a* decreases survival of KPC (*Pdx1-Cre, LSL-KRAS^G12D^, LSL-Trp53^R^*^172^*^H^*) mice and increases EGFR phosphorylation. (A) Kaplan-Meier survival curve of KPC PDAC mouse model showing decreased survival with loss of *Cip2a* (KPCC^fl/fl^, n=44) compared to wildtype (KPC, n=55) or heterozygous (KPCC^fl/+^, n=56). Significance was determined by log-rank Mantel-Cox (*p=*0.005) and Gehan-Breslow-Wilcoxon (*p=*0.002) tests. Mice sacrificed due to extra-pancreatic pathologies are censored and represented by ticks. (B) Representative images of endpoint tissue stained with PanCK to identify regions of interest and phosphorylated EGFR. Scale bars represent 100μm. (C) Quantification of mean fluorescent intensity of phospho-EGFR averaged from 5 fields of view from 6 tumors per genotype. *p<0.05 by two-tailed unpaired student’s t test

### Pharmacological PP2A activation functions synergistically with EGFR inhibition

EGFR signaling significantly contributes to the activation of oncogenic cascades that drive proliferation and survival in PDAC^24,25,45–47^. Therefore, it is possible that increased EGFR signaling in response to PP2A activation may attenuate the tumor suppressor functions of PP2A. In the current studies, we have determined that EGFR signaling is necessary for B56α-driven increase in PDAC cell proliferation. Therefore, we hypothesized that EGFR inhibitors may suppress the feedback loop between PP2A-B561? and EGFR, decreasing PDAC viability. To identify if the combination of PP2A activation and EGFR inhibition is efficacious, a drug matrix with DT-061 and the HER family inhibitor, Afatinib, was performed in 5 PDAC cell lines. SynergyFinder+ was used to calculate synergy between DT-061 and Afatinib and bliss synergy model was used^48^. The combination of DT-061 and Afatinib was synergistic in AsPc1, Panc1, and MiaPaCa2, but not in Capan2 or HPAFII (Figure 7A, B, Supplemental Figure 5A-C). We then tested if the combination of DT-061 and Afatinib was able to reduce tumor growth *in vivo*. Briefly, 2M MiaPaCa2 cells were injected subcutaneously into NOD.Cg-Rag1tm1Mom Il2rgtm1Wjl/SzJ (NRG mice). Once the average tumor size reached 100mm^3^, mice were randomly enrolled into four treatment arms: Vehicle, DT-061 (15mg/kg BID), Afatinib (15mg/kg QD), or Combination. Supporting our *in vitro* results, the combination of DT-061 and Afatinib significantly reduced tumor burden (Figure 7A) and endpoint tumor weight (Figure 7B), with no significant impact on mouse body weight (Supplemental Figure 5D). In whole cell tumor lysates, DT-061 treated tumors had a significant increase in phospho-Y1068 EGFR (pEGFR) expression levels compared to vehicle (Figure 7E). As expected, Afatinib effectively inhibited pEGFR, with no detectable phosphorylation at Y1068, and was able to prevent DT-061-driven pEGFR in combination with DT-061 (Figure 7E). These results indicate that inhibition of EGFR increases the tumor suppressive capabilities of PP2A.

**Figure 7.**
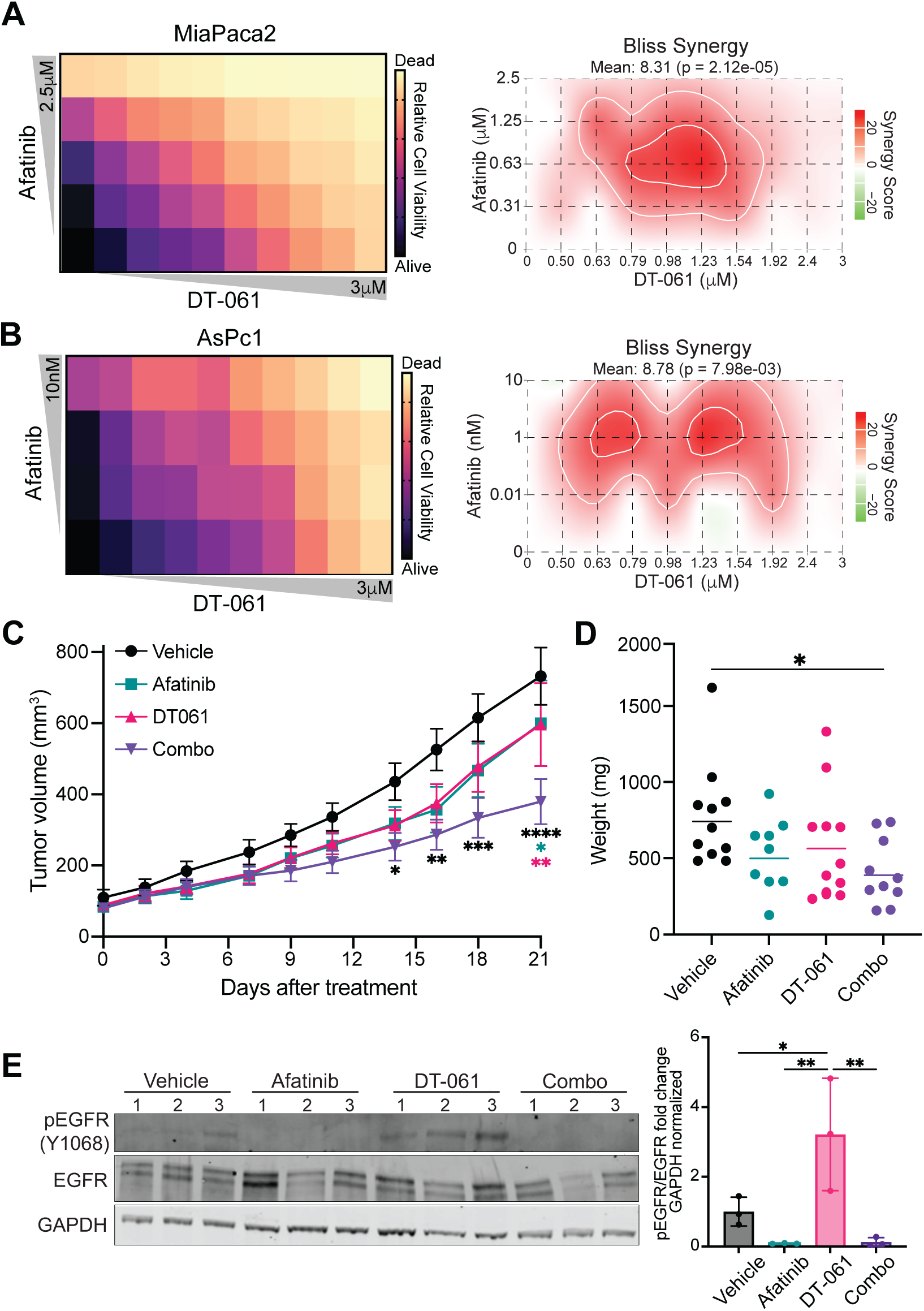
DT-061 and Afatinib treatment are a synergistic combination *in vitro* and *in vivo*. Drug matrix between Afatinib and DT-061 in (A) MiaPaCa2 and (B) AsPc1 cells. Average cell viability (left) was determined through crystal violet staining 72 hours after drug treatment from at least 3 independent biological replicates. Bliss synergy scores (right) across the matrix. (C) MiaPaCa2 cells were grown subcutaneously in NRG mice and mice were treated with vehicle (n=11), Afatinib (15mg/kg)(n=9), DT-061 (15mg/kg)(n=11), or combination (n=10) for 3 weeks. Mean ± SEM are shown. *p<0.05, **p<0.01, ***p<0.001, ****p<0.0001 by 2way ANOVA, significance of DT-061 + Afatinib compared to: vehicle (black stars), Afatinib (teal stars), DT-061 (pink stars) B by one-way ANOVA. (D) Final tumor weight from individual mice in panel C. *p<0.05 by 1-way ANOVA (E) Western blot from whole cell tumor lysates from individual tumors (n=3 tumors per genotype) from mice in panel C (left). Quantification of phosphorylated EGFR (Y1068) relative to total EGFR normalized to GAPDH (right). *p<0.05, **p<0.01 by 1-way ANOVA.

## DISCUSSION

PP2A represents a critical and complex regulator of cellular signaling cascades. Given the diverse functions of this large phosphatase family, a deep mechanistic understanding of the roles that PP2A plays during oncogenesis is currently lacking. Here, we have identified a novel mechanism by which activation of the specific PP2A subunit, B56α, results in increased signaling through the AREG-EGFR axis to drive proliferative phenotypes in PDAC. To our knowledge, these studies are the first to implicate B56α as functionally contributing to tumorigenic phenotypes and highlight the context specific functions of PP2A. Our findings also underscore the importance of defining patient populations that exhibit either tumor suppressive or tumor promotional phenotypes in response to PP2A signaling.

While PP2A is broadly considered to be tumor suppressive, examples of individual subunits exacerbating tumor phenotypes have been reported in several cancer cell types^49–52^ . B55α has been described as a tumor suppressive subunit in lung and breast cancer, but pro-tumorigenic in PDAC cell lines^53–56^. Although we were unable to recapitulate these findings with overexpression of B55α, differences in methods/approach or cell line could be a contributing factor. Furthermore, the context dependent functions of PP2A may vary across disease progression as our previous studies suggest that genetic loss of B56α accelerates pancreatic lesion formation during tumor initiation^57^. The increase in tumorigenic phenotypes in PDAC cell lines with B56α overexpression was not replicable in three KRAS-mutant NSCLC cell lines. While further studies are necessary in order to compare cancer specific phenotypes, these results support a potential differential role for B561Z in NSCLC and PDAC, and broadly suggest that different types of tumors or subsets of patients may display unique PP2A phenotypes. This concept becomes critically important in the context of therapeutic strategies that target PP2A.

Dynamic shifts in PP2A holoenzyme composition are poorly understood given the large number of B subunits and rapid cellular signaling responses, and most likely plays a significant role in determining the functional consequence of PP2A perturbations. There are also clear differences in biological responses between chronic and acute PP2A activation, with overexpression of B56α leading to increased proliferation (chronic adaptation to PP2A), while pharmacological activation of PP2A (acute activation) results in cell death. Given that our results for B56α overexpression and loss are complementary and coupled with pharmacological activation strategies give strength that B56α signaling alters EGFR pathways and tumorigenic phenotypes in PDAC. We also demonstrate that genetic loss of the PP2A inhibitor, CIP2A, leads to decreased survival *in vivo*. These results were surprising as we have previously demonstrated that loss of B56α accelerates the formation of pancreatic precursor lesions in a mutant KRAS model^57^ . CIP2A is known to regulate subunits in the B56 family; Therefore, this model likely represents the impact of multiple PP2A subunits beyond just B56α. Furthermore, CIP2A has PP2A-independent functions^58^. However, given that high CIP2A expression is often associated with aggressive tumor phenotypes, our findings suggest that the functional consequence of CIP2A-mediated regulation of PP2A is more complex than previously appreciated. Teasing apart the both the CIP2A and PP2A interactome during PDAC progression will provide critical insights as to how PP2A signaling changes during tumor progression and the impact of both high and low PP2A activity.

Finally, there was a significant increase in the amount of AREG within the media from DT-061 treated cells and B56αOE cell lines (Figure 4). Similar to AREG, the EGFR ligands EGF, HB-EGF, and EREG have all been implicated driving PDAC progression^59–64^ . However, we were unable to detect these ligands from the conditioned same media and, therefore, are unable to conclude if other ligands contribute to the activation EGFR downstream of PP2A. Given that knockdown of AREG attenuated the proliferation of B56αOE cells, AREG may be necessary for mediating this phenotype. AREG is a low affinity ligand and, therefore, may be available in the media for longer amounts of time^65,66^ . Furthermore, the fate of EGFR (internalization, recycling, degradation) downstream of AREG is poorly understood compared to ligands such as EGF . Future studies interrogating the dynamics of EGFR internalization and cellular fate in response to PP2A activation will shed light on the differential responses observed between cell lines and cell types. In addition to ligand-receptor dynamics, EGFR can function as a homodimer or heterodimer with other HER family proteins. As EGFR ligands have been shown to have differential impact on both the stabilization of EGFR dimers and EGFR binding partners, understanding the contribution of other HER family proteins to our phenotypes is critical^67^.

In summary, our studies suggest that cellular context plays a significant role in determining whether PP2A-B56α effectively functions as a tumor suppressor. In contrast to global PP2A functions, our findings identify EGFR signaling as a critical mediator of PDAC proliferation in response to PP2A-B56α activation and support therapeutic strategies to mitigate these phenotypes.

## ACKNOWLEDGEMENTS

We would like to thank all the members of the BAP lab for their contributions to this work, editing of the manuscript, and providing constructive feedback. The Cip2a^fl/fl^ mouse model was generated by Turku Center for Disease Modeling (www.tcdm.fi) supported by the Biocenter Finland. KPC survival curves generated in collaboration with the CRUK Scotland Institute’s Biological Services Unit. This study was supported by Purdue EVPRP NIH New R01 Program (BLA-P PI), Grace M. Showalter Research Trust (BLA-P); Pancreatic Cancer Action Network (BLA-P; 22-20-ALLE), Concern Foundation (BLA-P), and American Cancer Society (BLA-P; RSG-24-1259215-01-MM). BNH, ERDC, SJC, and EGS was supported by the Purdue Institute for Cancer Research (NIH grant P30 CA023168). JPM and SAK were supported by Cancer Research UK (A17196, A31287, A29996, A25233, CTRQQR-2021\100006). CMP was supported by the Purdue Bilsland Dissertation Fellowship. The bioinformatics analyses were supported by the Purdue Institute for Cancer Research and the Walther Cancer Foundation. The mass spectrometry work performed in this manuscript was done by the Indiana University School of Medicine Center for Proteome Analysis. Acquisition of the IUSM Center for Proteome Analysis instrumentation used for this project was provided by the Indiana University Precision Health Initiative. The proteomics work was supported, in part, by the Indiana Clinical and Translational Sciences Institute (Award Number UL1TR002529 from the National Institutes of Health, National Center for Advancing Translational Sciences, Clinical and Translational Sciences Award) and the P30 IU Simon Comprehensive Cancer Center Support Grant (Award Number P30CA082709 from the National Cancer Institute).

## AUTHOR CONTRIBUTION

CMP and BLA-P designed experiments. CMP, SJC, SAK, ERDC, BNH, LEG, EFK, GB, EGH, EGS, and EP performed and analyzed experiments. SMU, HK, and NAL performed RNA-seq data analysis. WS-K, KH, EHD, and ALM performed mass spectrometry sample preparation, data analysis, and methods Write-up. FZ and JW generated the Cip2a^fl/fl^ mouse. SAK and JPM generated KPC and KPCC survival curves and maintained the mouse colony. BLA-P secured funds and provided supervision. BLA-P and CMP wrote and revised the manuscript. All authors reviewed and finalized the manuscript.

## METHODS

### Cell culture

AsPc1, MiaPaCa2, HPAFII, Panc1, CfPac1, Capan1, Capan2, Colo357, and HEK293T were obtained from ATCC. PDAC cell lines were maintained in DMEM (Gibco 11995065) + 10% Fetal Bovine Serum (FBS) (Fisher Scientific FB12999102) at 37°C and 5% CO_2_. NSCLC cell lines (A549, H23, and Calu6) were maintained in RPMI (Cytiva SH30027.01) + 10% FBS at 37°C and 5% CO_2_. Lines were regularly tested for Mycoplasma using a PCR strategy. Viral particles were generated by transfecting HEK293T cells with the vector of interest and two lentiviral vectors (psPAX2 Addgene #12260 and pMD2.G Addgene #12259) using lipofectamine 3000 (Invitrogen L3000015). B56α knockout cell lines were generated by first generating Cas9 (Addgene #125592) expressing cells through clonal selection. Then the sgRNAs (sgNT: Horizon Discovery GSG11817; sgB56?1: Horizon Discovery GSGH11838-246527026; sgB56?2: Horizon Discovery GSGH11838-246527024) were transduced into the Cas9 expressing cells, selected and then clonal selected and screened for B56α loss by western blot. Overexpression of each of the B subunits were generated by cloning the gene of interest into a modified pSin-PURO empty vector. The following vectors were used for the B subunits: 4HA-B56α (Addgene #14532), 4HA-B5601 (Addgene #14534), 4HA-B56□ (Addgene #14536), 4HA-B56ε (Addgene #14537), Flag-B55α (Addgene #13804). For transient knockdown experiments, cells were transfected using lipofectamine 3000 with 35nM siRNA (NT (Horizon Discovery D-001810-20), EGFR (Horizon Discovery L-003114-00-005), AREG (Horizon Discovery L-017435-00-005), or B561? (Horizon Discovery L-009352-00-0020).

### Western blot

Cells were lysed and protein was quantified using DC Protein Assay Kit II (Bio-Rad 50000112). Equally loaded samples were separated on a 4-12% Criterion XT Bis-Tris Protein Gel (Bio-Rad 3450123) and transferred onto a PVDF membrane (Millipore IPFL00010). Membrane was blocked using Intercept (PBS) Blocking Buffer (Licor 927-70001) for 1 hour at room temperature. Primary antibodies were incubated overnight at 4°C and secondary antibodies were incubated at room temperature for 1 hour (Supplemental Table 9). Membranes were imaged with a LI-COR Odyssey and analyzed using Image Studio Lite.

Table of Antibodies

| Protein | Company | Catalog number | Lot number | Dilution |
| --- | --- | --- | --- | --- |
| Western Blot |  |  |  |  |
| EGFR | Cell Signaling | 2646S | 8 | 1:500 |
| p-EGFR (Y1068) | Cell Signaling | 2234L | 22 | 1:500 |
| AKT | Cell Signaling | 2920S | 8 | 1:1000 |
| p-AKT (S473) | Cell Signaling | 4058S | 30 | 1:1000 |
| HA | Cell Signaling | 3724S | 10 | 1:1000 |
| c-MYC | Abcam | AB32072 | 1053331-47 | 1:1000 |
| GAPDH | Invitrogen | AM4300 | 3035436 | 1:20,000 |
| B56α | Ogris lab | | | 1:500 |
| CIP2A | Santa Cruz | SC-80659 | H2522 | 1:500 |
| B55α | Cell Signaling | 5689T | 3 | 1:500 |
| IRDye 680RD Goat Anti-Rabbit IgG Secondary | Licor | 926-68071 | D20809-05 | 1:5000 |
| IRDye 800CW Goat Anti-Mouse IgG Secondary | Licor | 926-32210 | D20802-25 | 1:5000 |
| Immunofluorescence |  |  |  |  |
| p-EGFR (Y1068) | Cell Signaling | 2234L | 22 | 1:100 |
| Pan Cytokeratin C11 | Santa Cruz | sc-8018 | J0721 | 1:1000 |
| Goat anti-Rabbit IgG (H+L) Alexa Fluor Plus 647 | Invitrogen | A32733 | ZJ405179 | 1:200 |
| Goat anti-Mouse IgG (H+L) Alexa Fluor Plus 488 | Invitrogen | A32723 | ZH395844 | 1:200 |

### RNA isolation/quantitative PCR

RNA was isolated using GeneJet RNA Purification Kit (ThermoScientific K0732) according to manufacturer’s protocol. cDNA was generated using the High-Capacity cDNA Reverse Transcription Kit (Applied Biosystems 4368814). Quantitative PCR was run using PowerUp SYBR Green Master Mix (Applied Biosystems A25741) using primers (Supplemental Table 10) on a QuantStudio3. Fold change was calculated using the delta delta ct method and normalized to 18S.

Table of Primers

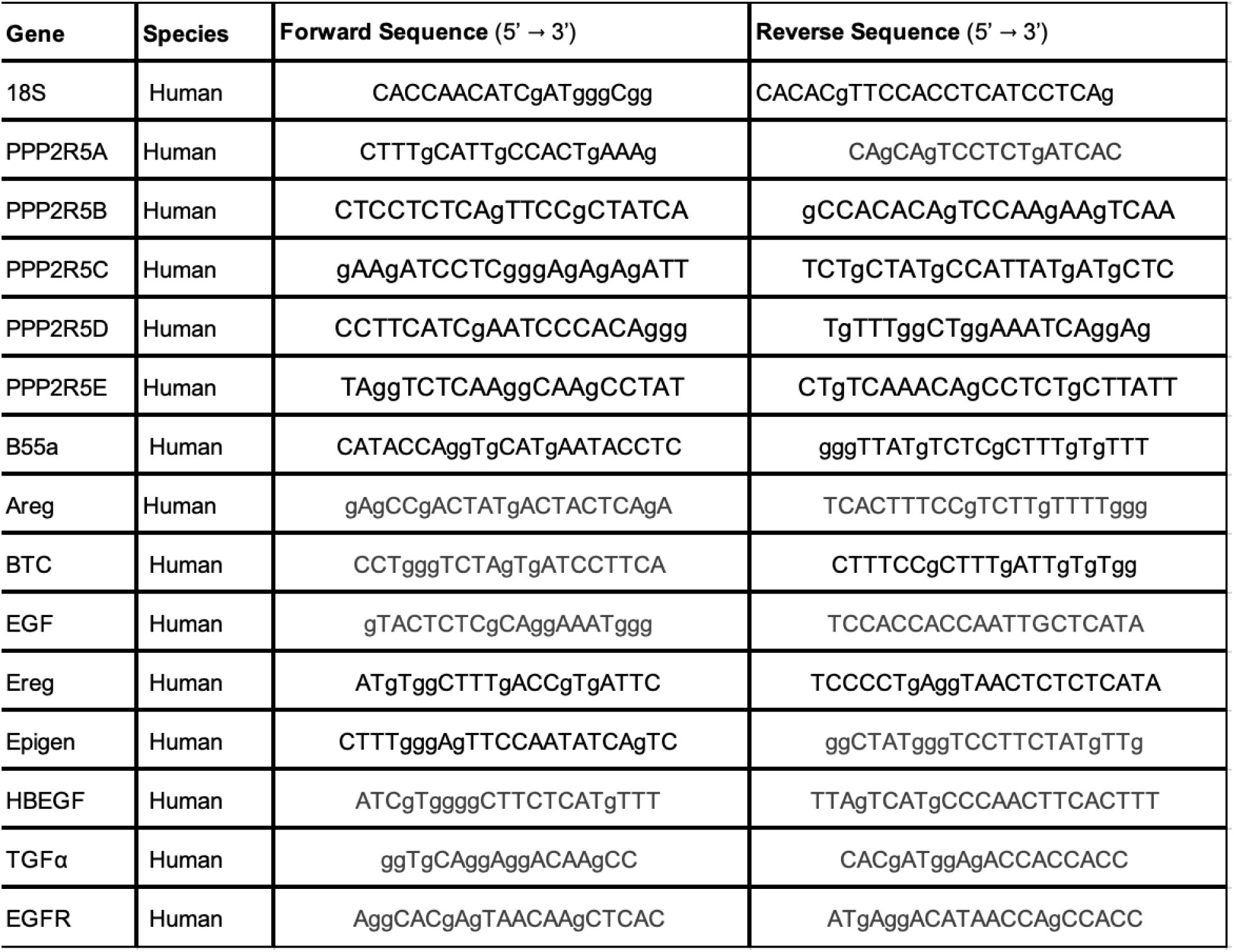

### Tumorigenic phenotypes

For proliferation, cells were seeded in technical triplicate and counted every 24 hours. Area under the curve was calculated and used for statistical significance. For soft agar assays, 12 well plates were coated with 500μL of a 1:1 mixture of 1.4% noble agar and 2X DMEM + 20% FBS. Based on the parental cell line, 30,000-50,000 cells were resuspended in a 1:1 mixture of 0.7% noble agar and 2X DMEM + 20% FBS and seeded on top of the 1.4% layer. After 14 days, cells were stained with 0.1% crystal violet and imaged using an EVOS M7000. Number of colonies was quantified using ImageJ. For migration, transwell migration assay was used. 1 hour prior to seeding, 8μm pore inserts (Corning 3464) were soaked in serum-free DMEM. 20,000 cells were seeded in 8μm inserts in serum-free DMEM and DMEM + 10% FBS was added to the bottom of the well. After 48 hours, cells were fixed and stained with 0.1% crystal violet. Cells that remained at the top of the insert and had not migrated were removed. Migrated cells were imaged with an EVOS M7000 and number of cells migrated was quantified using ImageJ. For clonogenic colony formation, 100-500 cells (depending on the parental cell line) were seeded in a 6-well plate. After 10-14 days (depending on parental cell line), cells were stained with 0.1% crystal violet and scanned on an EVOS M7000. Percent area occupied by colonies was quantified using ImageJ.

### RNA sequencing

AsPc1, Panc1, and HPAFII cells were plated in DMEM +2% FBS in triplicate. 48 hours later, cells were treated with 10μM DT-061 or vehicle (DMSO) for 24 hours. RNA was isolated as described above. RNA-seq libraries were generated and Illumina sequenced by Novogene. Datasets were trimmed using fastp (v0.23.2) to remove bases with a Phred quality score lower than Q30 and adaptor sequences. Reads were aligned to GRCh38.p14 using STAR version 2.7.9a^68^, and genes were quantified with subread featureCounts version 1.6.1^69^ using the annotation from ensembl release 104. Initial exploratory analyses were performed with DESeq2 version 1.26.0^70^ and differential gene expression analyses were performed with edgeR version 3.28.1^71^ on the R Statistical Computing Project version 3.6.1^72^ . Ranked lists were generated for each cell line and used for Gene Set Enrichment Analysis (GSEA) version 4.3.2.^22,23^.

### Total and phospho proteomics

5x10^6^ AsPc1 cells were plated onto a 15cm plate in DMEM + 2% FBS. 48 hours later, cells were treated with 5μM DT-061 or vehicle for 24 hours. Cells were lifted, washed, and then pelleted and flash frozen. Sample preparation, mass spectrometry analysis, bioinformatics, and data evaluation for quantitative proteomics experiments were performed in collaboration with the Indiana University Proteomics Center for Proteome Analysis at the Indiana University School of Medicine similarly to previously published protocols^73,74^.

#### Protein extraction and digestion

Cell pellets were resuspended in 100 µL 8 M Urea, 100 mM Tris pH 8.5. The resuspended cell pellets were transferred to Diagenode Bioruptor tubes (Cat No: C30010010). Cells were lysed via Diagenode Bioruptor, 30s on/30s off, for 30 cycles. Samples were incubated for 1 hour at 35 °C with 2 units of benzonase (EMD Millipore Cat NO. 70746-4). Samples were then clarified by centrifuging for 20 min at 12,000 rcf. Supernatants were analyzed in a Bradford assay (Biorad Cat No: 5000002) to determine protein concentration. 520 μg of each sample was treated with 5 mM tris(2-carboxyethyl)phosphine hydrochloride (Sigma-Aldrich Cat No: C4706) to reduce disulfide bonds and the resulting free cysteine thiols were alkylated with 10 mM chloroacetamide (Sigma Aldrich Cat No: C0267). Samples were diluted with 50 mM Tris.HCl pH 8.5 (Sigma-Aldrich Cat No: 10812846001) to a final urea concentration of 2 M for overnight Trypsin Platinum digestion at 35 °C (1:25 protease:substrate ratio, Mass Spectrometry grade, Promega Corporation, Cat No: VA9000).

#### Peptide purification and phosphopeptide enrichment

Digestions were quenched with formic acid (Fluka Cat No: 94318) and desalted on Waters Sep-Pak® Vac cartridges (Waters™ Cat No: WAT054955) with a wash of 1 mL 0.1% TFA followed by elution in 3x 0.2 mL of 70% acetonitrile 0.1% formic acid (FA). Peptides were dried by speed vacuum. Samples were resuspended in phosphopeptide binding buffer and phosphopeptides were enriched using Thermo Fisher Scientific High Select TiO2 tips (Cat No A32993) according to manufacturer’s instructions. Flow through (non-phosphopeptides) and phosphopeptides were dried down by speed vacuum.

#### TMTpro labeling

An equivalent of 50μg of the global (non-phospho) peptides and all of each phosphopeptide enrichment were resuspended in 100 mM triethylammonium bicarbonate (TEAB, pH 8.5 from 1 M stock). Each sample was then labeled overnight at room temperature, with 0.5 mg of Tandem Mass Tag Pro (TMTpro™) reagent (16-plex kit, manufactures instructions Thermo Fisher Scientific, TMTpro™ Isobaric Label Reagent Set; Cat No: 44520, lot no. ZA382395. Reactions were quenched with 0.3 % hydroxylamine (v/v) at room temperature for 10 minutes. Labeled peptides were then mixed and dried by speed vacuum.

#### High pH basic fractionation

Half of the combined global sample and all of the phosphopeptide sample were resuspended in 0.5% TFA and fractionated on a Waters Sep-Pak® Vac cartridge (Waters™ Cat No: WAT054955) with a 1 mL wash of water, 1 mL wash of 5% acetonitrile, 0.1% triethylamine (TEA) followed by elution for the global sample in 8 fractions of 12.5%, 15%, 17.5%, 20%, 22.5%, 25%, 30%, and 70% acetonitrile, all with 0.1% TEA).

#### Nano-LC-MS/MS

Mass spectrometry was performed utilizing an EASY-nLC 1200 HPLC system (SCR: 014993, Thermo Fisher Scientific) coupled to Eclipse™ mass spectrometer with FAIMSpro interface (Thermo Fisher Scientific). Each multiplex was run on a 25 cm Aurora Ultimate column (Ion Opticks Cat No: AUR3-25075C18) in a 50 °C column oven with a 180-minute gradient or 130-minute gradient. For each global fraction, 6.67% of the sample was loaded and run at 350 nl/min with a gradient of 10-40%B over 160 minutes; 40-90% B over 10 mins; held at 90% for 2 minutes; and dropping from 90-8% B over the final 5 min (Mobile phases A: 0.1% formic acid (FA), water; B: 0.1% FA, 80% Acetonitrile (Thermo Fisher Scientific Cat No: LS122500)). The mass spectrometer was operated in positive ion mode, default charge state of 2, advanced peak determination on, and EasyIC on. Three FAIMS CVs were utilized (-45 CV; -55 CV; -65CV) each with a cycle time of 2 s and with identical MS and MS2 parameters. Precursor scans (m/z 400-1600) were done with an orbitrap resolution of 120000, RF lens% 30, 50 ms maximum inject time, standard automatic gain control (AGC) target, minimum MS2 intensity threshold of 2.5e4, MIPS mode to peptide, including charges of 2 to 6 for fragmentation with 60 sec dynamic exclusion shared across the cycles excluding isotopes. MS2 scans were performed with a quadrupole isolation window of 0.7 m/z, 34% HCD collision energy, 50000 resolution, 200% AGC target, dynamic maximum IT, fixed first mass of 100 m/z.

For each phosphopeptide fraction 25% of the sample was loaded and run at 350 nl/min with a gradient of 10-35%B over 120 minutes; 35-95% B over 5 mins; and dropping from 95-6% B over the final 5 min (Mobile phases A: 0.1% formic acid (FA), water; B: 0.1% FA, 80% Acetonitrile (Thermo Fisher Scientific Cat No: LS122500)) with identical mass spectrometry parameters to the global proteomics samples.

#### Mass spectrometry data analysis

Resulting RAW files were analyzed in Proteome Discover™ 2.5.0.400 (Thermo Fisher Scientific)^75^ with a *Mus musculus* UniProt reference proteome FASTA (downloaded 051322) plus common laboratory contaminants (73 sequences). SEQUEST HT searches were conducted with full trypsin digest, 3 maximum number missed cleavages; precursor mass tolerance of 10 ppm; and a fragment mass tolerance of 0.02 Da. Static modifications used for the search were: 1) TMTpro label on peptide N-termini 2) TMTpro label on lysine (K) and 3) carbamidomethylation on cysteine (C) residues. Dynamic modifications used for the search were 1) oxidation on M (2) phosphorylation on S, T or Y, 3) acetylation on protein N-termini, 4) methionine loss on protein N-termini or 5) acetylation with methionine loss on protein N-termini. A maximum of 3 dynamic modifications were allowed per peptide. Percolator False Discovery Rate was set to a strict setting of 0.01 and a relaxed setting of 0.05. IMP-ptm-RS node was used for all modification site localization scores. Values from both unique and razor peptides were used for quantification. In the consensus workflows, peptides were normalized by total peptide amount with no scaling. Unique and razor peptides were used, and all peptides were used for protein normalization and roll-up. Quantification methods utilized TMTpro isotopic impurity levels available from Thermo Fisher Scientific. Reporter ion quantification filters were set to an average S/N threshold of 5 and co-isolation threshold of 30%. Resulting grouped abundance values for each sample type, abundance ratio values; and respective p-values (Protein Abundance based with ANOVA individual protein based) from Proteome Discover were exported to Microsoft Excel. Pathway enrichment was performed using Gene Set Enrichment Analysis (GSEA) version 4.3.2^21–23^.

### Conditioned media

2.5M AsPc1 B561ZOE or EV cells were plated in a 10cm plate in DMEM +10% FBS. 48 hours later, media was changed to 5.5ml serum free DMEM and allowed to condition for 48 hours. Conditioned media was filtered through a 0.45μM filter and stored at 4°C until use. 2ml of conditioned media was added to AsPc1 parental cells 48 hours after seeding 200k on 35mm plates. After 90 minutes, cells receiving conditioned media were lysed and protein was analyzed as described in western blotting section. The same process was used with DT-061 treated cells with the following modifications: 2.5M AsPc1 cells were plated in a 10cm plate in DMEM + 10% FBS. 24 hours later, media was changed to serum free DMEM and 7.5μM DT-061 was added for 6 hours, then drug was washed out and serum free DMEM was allowed to condition for 48 hours. After 24 hours, cells receiving conditioned media were lysed and protein was analyzed as described in western blotting section.

### ELISA

To collect media, 300,000 cells were seeded in a 6-well plate in DMEM + 10% FBS. 24 hours after seeding, cells were serum starved for 24 hours prior to adding DMEM to condition for 24 hours. Media was stored at -20°C until quantified for AREG (Invitrogen EHAREG), HB-EGF (Invitrogen EHHBEGF), EGF (Invitrogen KHG0061), or EREG (Abcam ab277077) according to manufacturer’s instructions. For the KPC/KPCC cell lines, AREG (Invitrogen EMAREG) was quantified and normalized to cell number for each biological replicate.

### B56α Rescue experiments

2M AsPc1 EV or B561ZOE cells were seeded and 24 hours later, lipofectamine 3000 (Invitrogen, L3000015) was used to transfect 35nM siRNA (NT (Horizon Discovery D-001810-10-20), EGFR (Horizon Discovery L-003114-00-005), or AREG (Horizon Discovery L-017435-00-005)). After 24 hours later, the cells were lifted, and the rescue experiments (proliferation and migration) were plated as described above. Cells were taken at all time points to validate knockdown of target gene via qPCR.

### Drug curves and matrices

7,000 cells were seeded and immediately dosed with Afatinib and DT-061. 72 hours later, cell viability was assessed using crystal violet^76^. Absorbance (570nm) was read on a Synergy H1 Microplate Reader (BioTek). Viability was calculated relative to vehicle control. 4 biological replicates were averaged, and synergy was calculated using the bliss synergy model in SynergyFinder+^48^.

### Animal studies

Animal experiments were performed in compliance with Purdue University Institutional Animal Care and Use Committee, University of Glasgow Animal Welfare and Ethics Review Board under UK Home Office project license (PP8411096), and the Finnish ethical committee for experimental animals, complying with international guidelines on the care and use of laboratory animals.

*Xenograft Studies:* MiaPaCa2 cells (2x10^6^) were injected subcutaneously into the flanks of NRG (NOD-*Rag1*^null^ *IL2Rg*^null^) mice in a 1:1 ratio with Matrigel (Corning 356231). Once palpable, tumors were calipered every other day and tumor volume was calculated by Volume = (length x width^2^)/2. Once average tumor volume was 100mm^3^, mice were randomly enrolled into the four treatment arms. 15mg/kg DT-061 or vehicle (1:1:8 N,N dimethylacetamide (DMA): Kolliphor HS 15: PBS) were dosed two times daily via oral gavage. 15mg/kg Afatinib or vehicle (1:1:18 N,N dimethylacetamide (DMA): Kolliphor HS 15: PBS) were dosed once daily by intraperitoneal injection. Mice were treated for 21 days and then mice were sacrificed. Tumors were halved and fixed in 10% formalin or snap frozen for qPCR and western blot. For histology, tumors were paraffin embedded and 6um sections were used H&E and immunofluorescence.

*Cip2a^fl/fl^ mice*: Generation of *Cip2a^fl/fl^* mice was done in collaboration with the Turku Center for Disease Modeling (TCDM) at the University of Turku (Finland). LoxP sites were inserted into introns 1 and 2 using the following strategy: BAC clones containing mouse *Cip2a* gene (ENSMUSG00000033031) were obtained from BACPAC Resources Center (Children’s Hospital Oakland Research Institute, Oakland (CHORI), California, USA). A 9696 bp genomic DNA fragment containing exons 1 to 8 of Cip2a gene was subcloned into the minimal vector, pACYC177 (New England Biolabs, MA, USA). To insert a single loxP site, a floxed PGK-Neo cassette (Gene Bridges) containing 50 bp *Cip2a* homology arms amplified by PCR was inserted into the intron 1 of *Cip2a* gene by Red/ET recombination method. Primers were 5’-TAGAGAAGGAAGAACTCTATATATGGGCAGAATGAACAAAAGATGTGAATGAAGCAGGGATTCTGCAAAC-3’ and 5’-AAACCAATTTTCTAAATTGCTACTTAGCCCTTTCTTTCCCAGTTTAGAAAGGCGGATTTGTCCTACTCAGG-3’. Site-specific recombination was carried out *in vitro* using Cre recombinase in E.coli. (Gene Bridges). A second LoxP site was introduced into the intron 2 of *Cip2a* gene with Frt-PGK-gb2-Neo-Frt-loxP cassette (Gene Bridges) containing 50 bp *Cip2a* homology arms amplified by PCR. The primers were 5’-ATAGATAGAATTATAAAATTTCTTCAACAATGATTTACATTCAAAAGATTAATTAACCCTCACTAAAGGGCGG-3’ and 5’-GTAGGAACTTTGAGGTATCTGTAATACTAACTCTAAAAGGTTTTACAATATAATACGACTCACTATAGGG-3’. G4 embryonic stem cells (derived from mouse 129S6/C57BL/6Ncr) were electroporated with 30 μg of linearized targeting construct. In order to delete Neo cassette in the targeted ES cells, cells were re-electroporated with plasmid, pCAGGS-Flpe. Targeted ES cells with Neo deletion were then injected into C57BL/6 (Charles River Laboratories) mouse blastocysts to generate chimeric mice. Germline transmission was achieved by cross-breeding male chimeras with C57BL/6 females. *Cip2a^fl/fl^* mice were maintained in a specific pathogen free stage at Central Animal Laboratory at the University of Turku before being transferred to the CRUK Scotland Institute. To generate KPCC mice, *Cip2a^fl/fl^* were bred with *Pdx1-Cre; LSL-Kras^G12D^; LSL-Trp53^R^*^172^*^H^*mice. Mice were maintained on a mixed background in conventional cages with environmental enrichment and given access to standard diet and water ad libitum. Mice of both sexes, in similar proportions, were used in all cohorts. Genotyping was performed by Transnetyx (Cordoba, TN, USA). Both KPC and KPCC mice were monitored at least three times per week by researchers blinded to genotype and humanely sacrificed when exhibiting moderate clinical signs of PDAC (swollen abdomen, loss of body conditioning, lethargy).

### Immunofluorescence

Paraffin embedded tissue slides were incubated at 60°^C^ for 1 hour prior to progressive rehydration in xylene twice for 10 minutes, then 100% ethanol twice for 5 minutes, then 95% ethanol, 70% ethanol, PBS, and water for 5 minutes each. Antigen retrieval was performed with slides first covered with 1x high pH antigen retrieval solution (Invitrogen 00-4956-58) and cooked in a pressure cooker on high pressure for 10 minutes. The slides were then allowed to cool to room temperature and rinsed in water before being covered with 1x low pH antigen retrieval solution (Invitrogen 00-4955-58) and cooked as described above. After cooling to room temperature, the tissues were covered with 0.025% tween in PBS for five minutes. Each tissue was blocked with 3% BSA in PBS for 1 hour at room temperature before the addition of primary antibodies diluted in 3% BSA in PBS and incubation overnight at 4°C. Tissues underwent three 5-minute washes with 0.025% tween in PBS before the addition of secondary antibodies which were incubated at room temperature for 1 hour. Secondary antibodies diluted in 3% BSA in PBS were followed with another series of three 5-minute washes in 0.025% tween in PBS. Dilutions of antibodies used are listed in (Supplemental Table 9). DAPI was diluted in 3% BSA in PBS before incubation on the tissues for 5 minutes at room temperature. After an additional 5-minute wash in 0.025% tween in PBS, a glass coverslip was mounted to each slide with Prolong Gold (Invitrogen P36934). Coverslips were allowed to harden for at least 12 hours before microscopy. A Nikon Ni-U Upright Microscope was used to take 20x images of each tissue. From each tissue, mean fluorescence intensity of phospho-EGFR (pY1068) from four to eight fields of view was quantified using ImageJ.

### Statistical Analysis

Technical replicate is defined as duplicate conditions within a single experiment to assess technical variability. Biological replicate is defined as experiments that are independently repeated in order to assess biological variability. All experiments have been performed with three independent biological replicates, with each individual experiment having at least two replicate conditions unless otherwise noted. Error bars represent standard deviation unless otherwise noted. For two groups, a student’s two-tailed unpaired t-test was performed using GraphPad Prism with 95% confidence interval. For groups of three or more, a 1-way ANOVA

