## Supplemental Figures for "Activation of PP2A-B56α leads to aberrant EGFR signaling and proliferative phenotypes in PDAC"

### Supplemental Figure 1

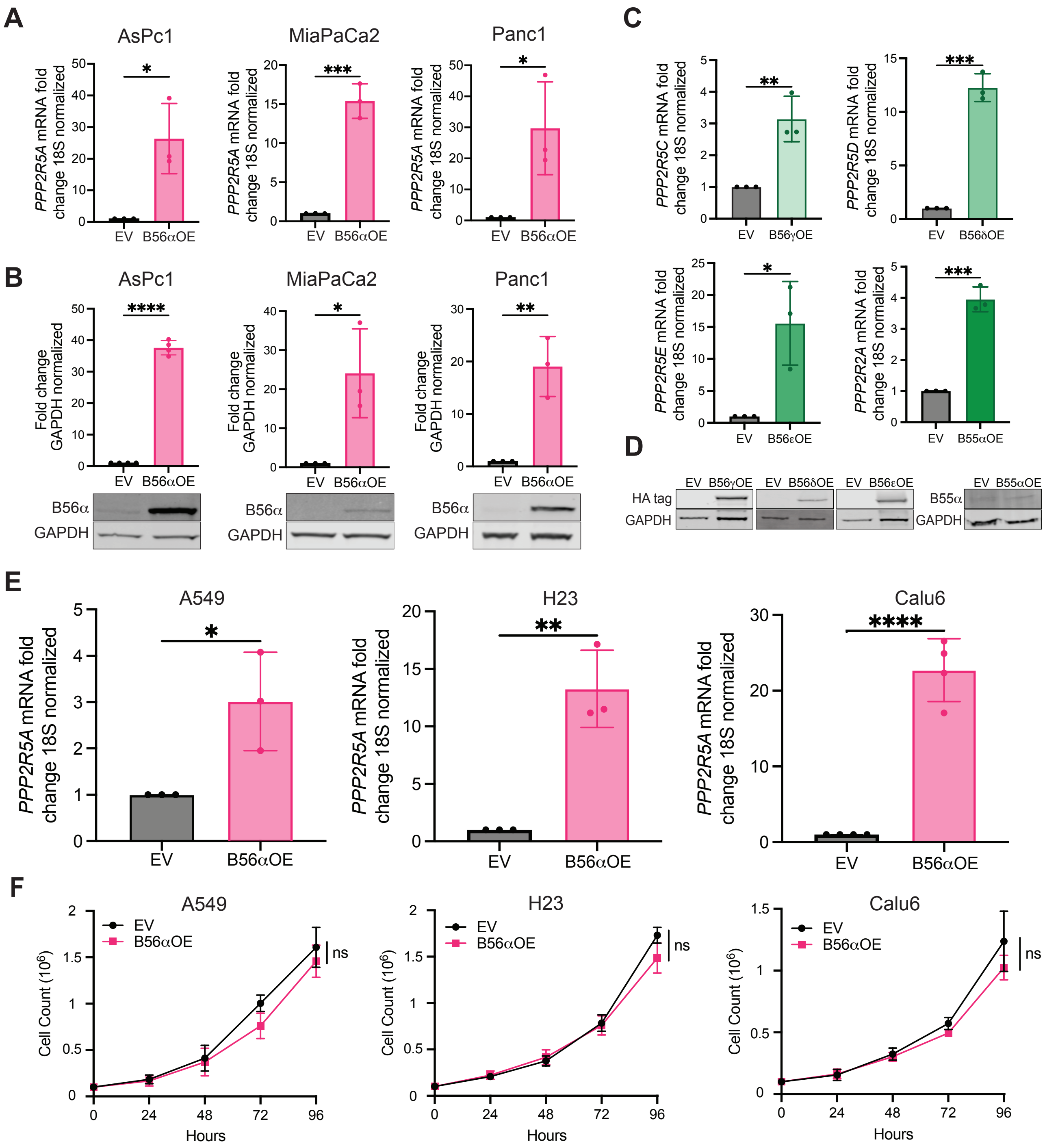

### Supplemental Figure 2

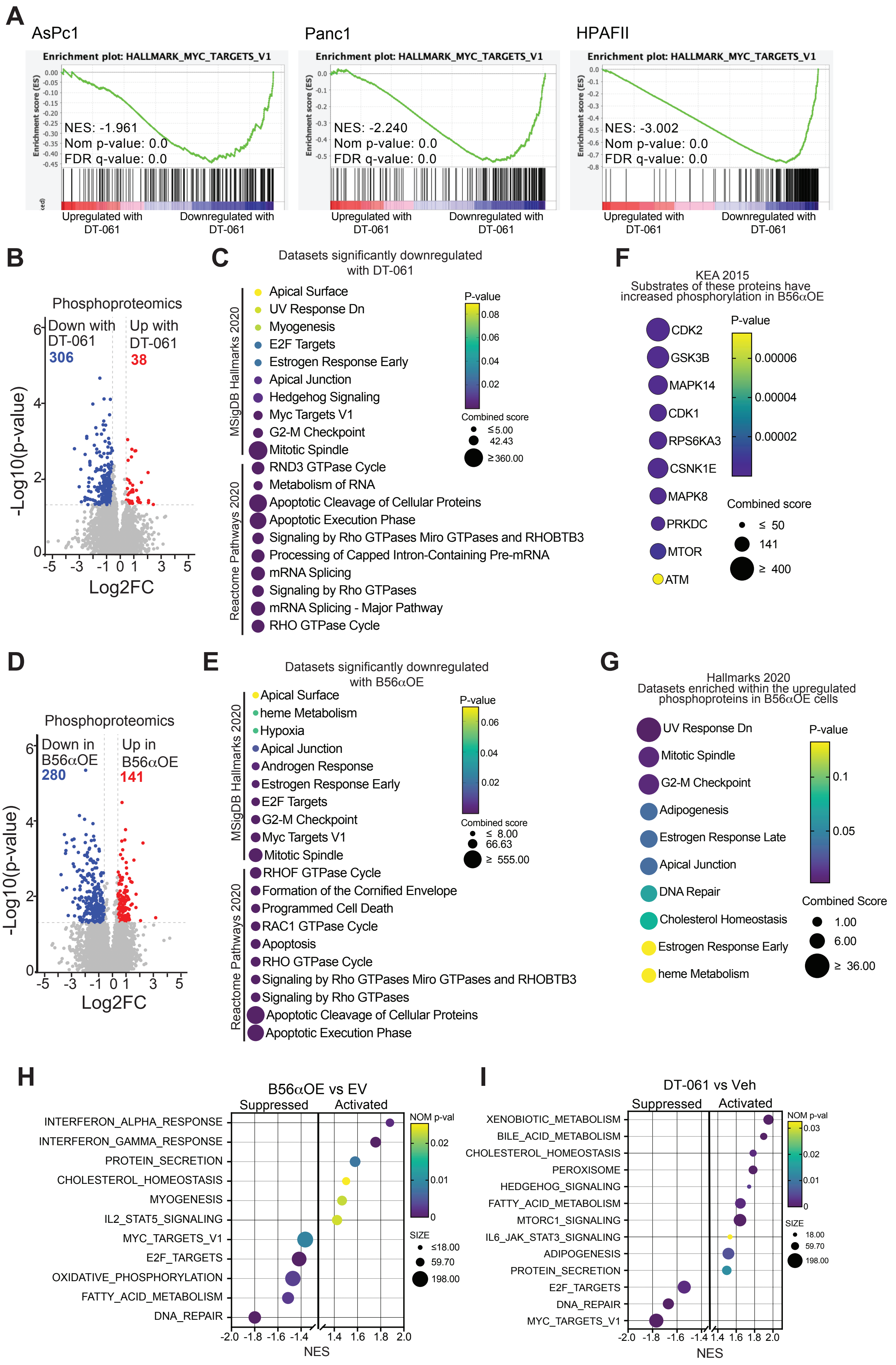

### Supplemental Figure 3

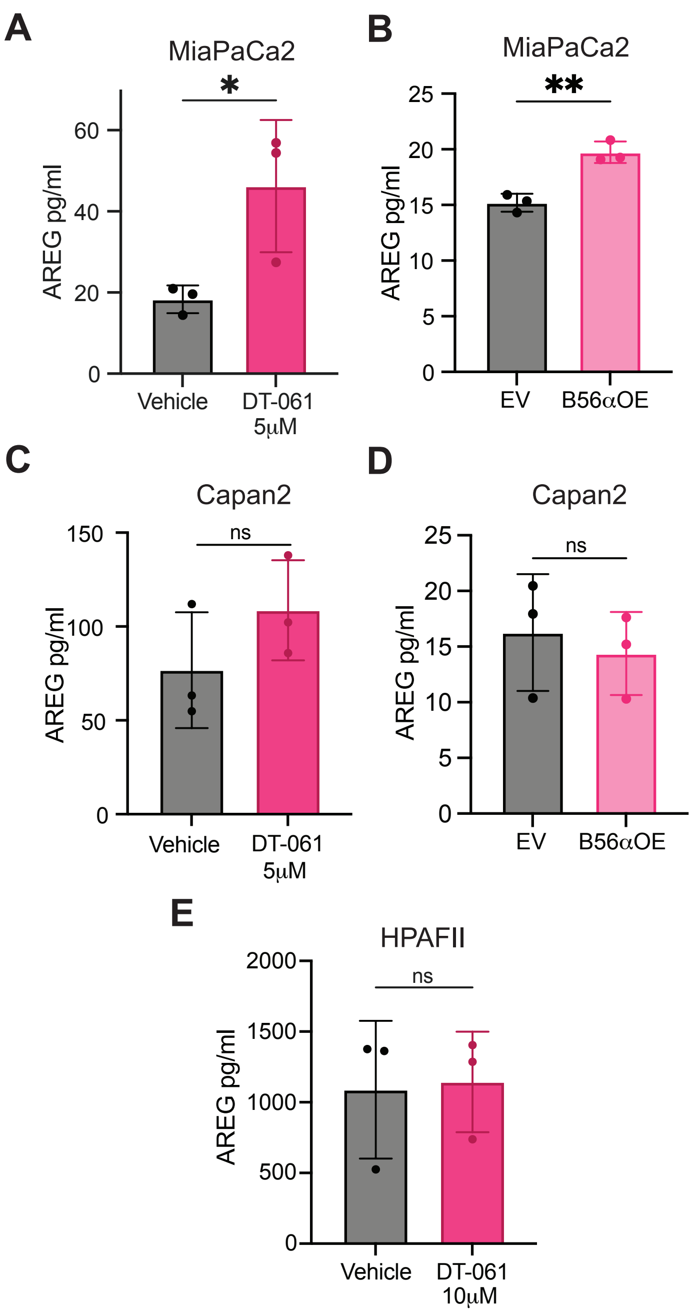

### Supplemental Figure 4

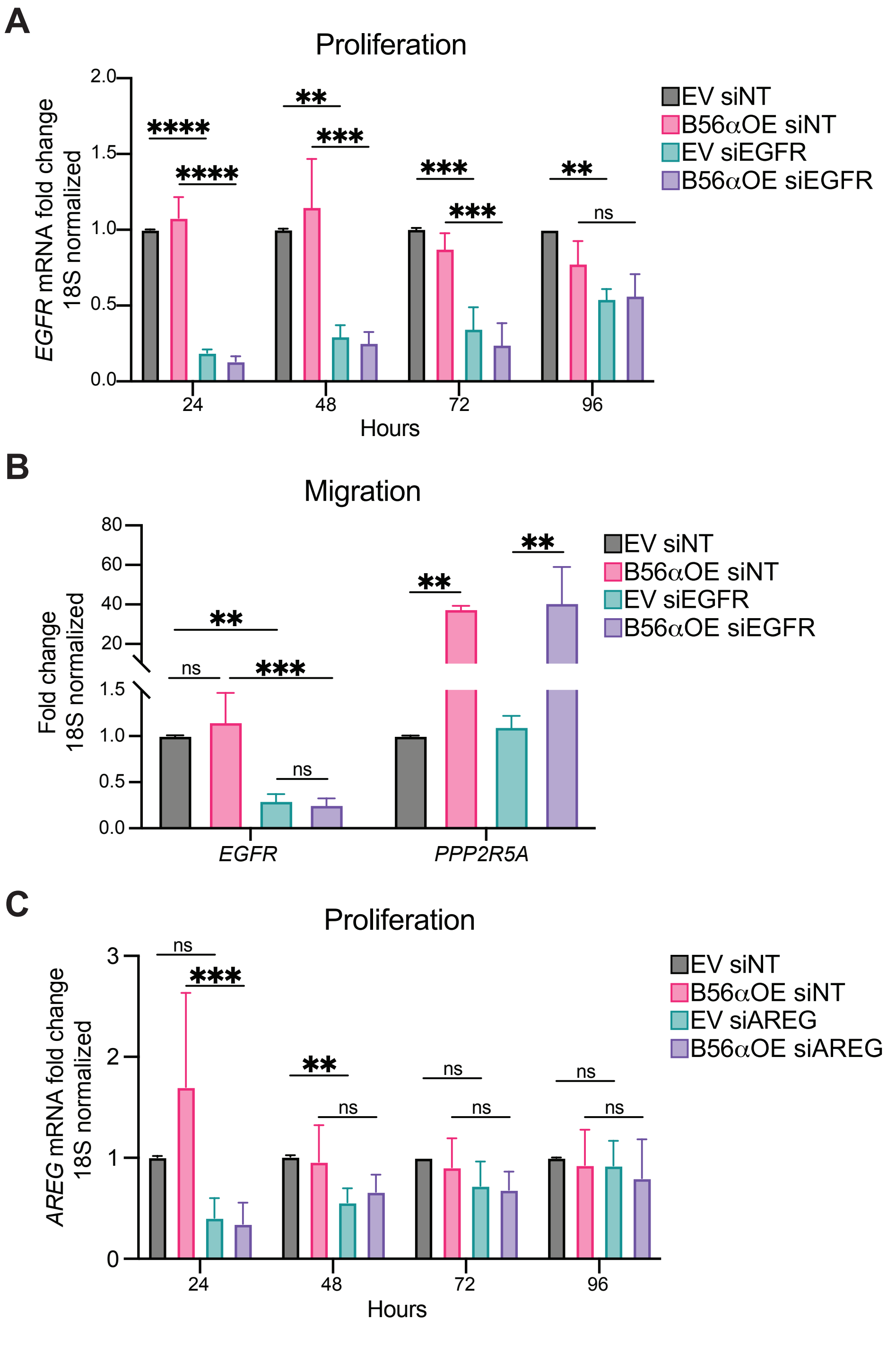

### Supplemental Figure 5

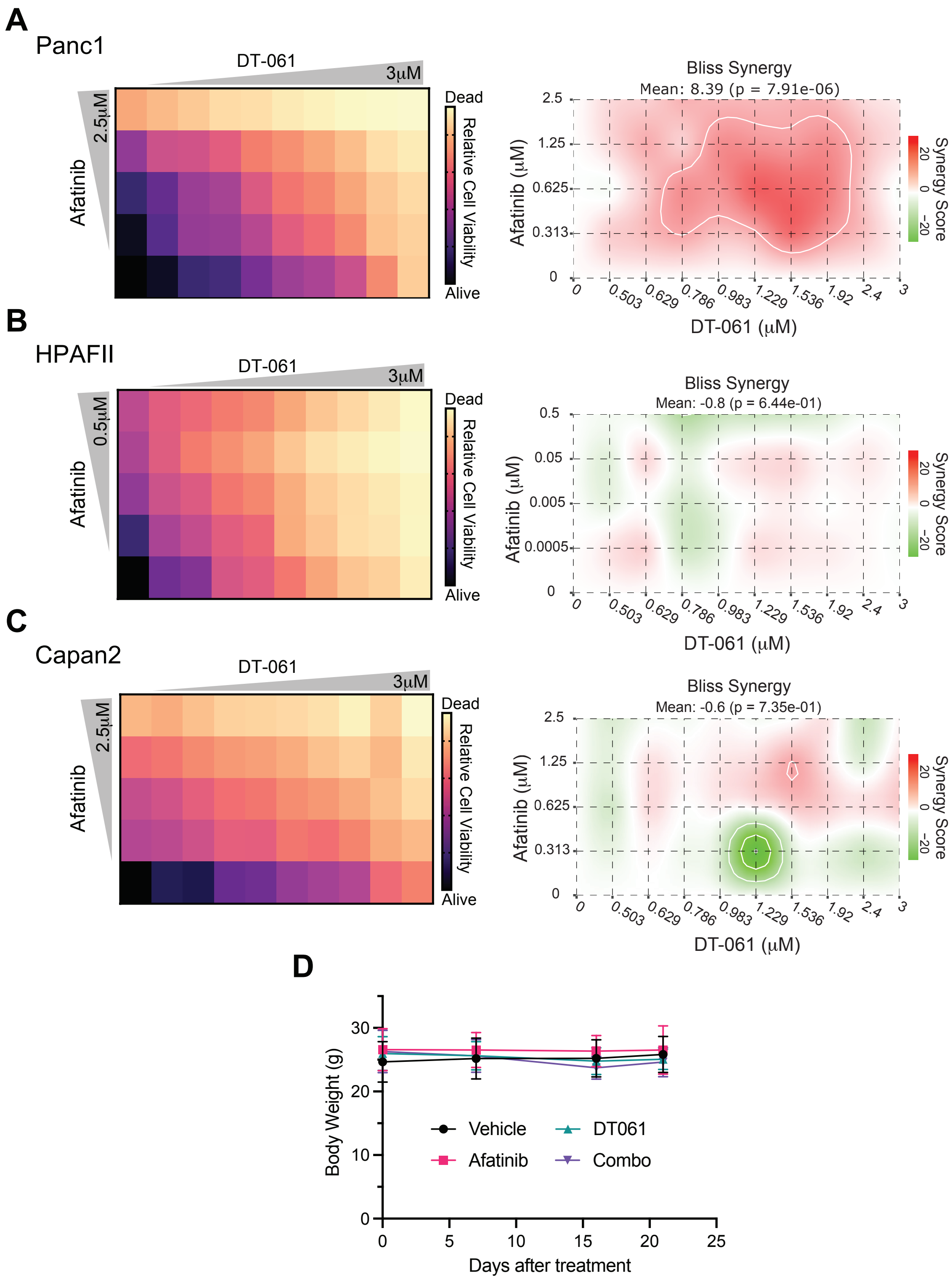
